# Differential Histone-DNA Interactions Dictate Nucleosome Recognition of the Pioneer Transcription Factor Sox

**DOI:** 10.1101/2021.12.07.471700

**Authors:** Burcu Ozden, Ramachandran Boopathi, Ayşe Berçin Barlas, Imtiaz N. Lone, Jan Bednar, Carlo Petosa, Seyit Kale, Ali Hamiche, Dimitar Angelov, Stefan Dimitrov, Ezgi Karaca

## Abstract

Pioneer transcription factors (PTFs) have the remarkable ability to directly bind to chromatin for stimulating vital cellular processes. In this work, we dissect the universal binding mode of Sox PTF by combining extensive molecular simulations and DNA footprinting techniques. As a result, we show that when Sox consensus DNA is located at the solvent-facing DNA strand, Sox binds to the compact nucleosome without imposing any significant conformational changes. We also reveal that the base-specific Sox:DNA interactions (base reading) and the Sox-induced DNA changes (shape reading) are concurrently required for the sequence-specific DNA recognition. Among different nucleosomal positions, such a specific reading mechanism is satisfied solely at superhelical location 2 (SHL2). While SHL2 acts transparently to Sox binding, SHL4 permits only shape reading, and SHL0 (dyad) allows no reading mechanism. These findings demonstrate for the first time that Sox-based nucleosome recognition is essentially guided by the distinct histone-DNA interactions, permitting varying degrees of DNA flexibility.

## INTRODUCTION

The nucleosome core particle (NCP) is the basic repeating unit of the eukaryotic genome (1). The NCPs, connected with the linker DNA, make up the 10 nm chromatin filament, which folds into higher order chromatin structures upon binding to the linker histone H1 (2–5). The NCP comprises of a core histone octamer (twice the H2A, H2B, H3, and H4 proteins) and 147 base pairs (bp) of DNA. The 147 bp DNA wraps around the histone octamer in 1.67 left-handed helical turns (6), of which positioning on NCP is described by the superhelical locations (SHLs). The SHLs are counted in the positive and negative directions, starting from the central DNA sequence, i.e., the nucleosomal dyad (SHL0). They are separated by ~10 bp nucleotides while running from ± 7 to ±1 positions (7). At each SHL, the accessibility of DNA for proteins is regulated by chromatin remodelers (8). The remodelers aid the freeing of nucleosomal DNA from histones to make specific recognition sequences available for transcription factors (TFs) (9, 10). In the absence of remodelers, nucleosomes present an impediment to transcription, as most TFs cannot overcome the nucleosomal barrier to recognize their binding sequence (11–17). An exception to this rule is the pioneer transcription factors (PTFs). Unlike conventional TFs, PTFs directly bind to nucleosomally organized DNA to assist the assembly of complex transcriptional machineries (18–20). This fact delegates PTFs a central role in essential chromatin-templated processes, such as establishing competence for gene expression and initiating cellular programming (21).

During the past decade, several studies explored the interaction landscape between PTFs and the NCP (18–20). According to what was observed, PTFs are proposed to have HMG, Fork-head, POU, ZF, and bHLH domains (20, 22). Among these domains, the HMG is composed of 79 amino acids, folding into a simple three-helix architecture arranged in a boomerang L-shape (23–26) (Figure 1A). Like the other PTFs, HMGs are demonstrated to guide a diverse range of vital processes (27–34). For example, the HMG domain carrying Sox2 was shown to be a part of the TF cocktail, capable of inducing pluripotent stem cells from somatic human cells (35, 36). Another HMG protein, Sox4, is a crucial factor in the epithelial to mesenchymal transition, a fundamental process operating in cancer progression and metastasis (37, 38). Together with the other eighteen proteins, Sox2/4 make up the Sox family, the so-called *“ultimate utility player of the cell”* (36, 39). For the sake of simplicity, from this point and on, we will refer to Sox-HMG as Sox. The Sox proteins recognize the cognate 5’-TTGT-3’ sequence, positioned at a DNA minor groove upon establishing base-specific hydrogen bonds (base reading), as well as upon imposing extreme DNA distortion induced by the hydrophobic Sox:DNA interactions (shape reading) (39–42) (Figure 1A). These reading mechanisms are invariant among all Sox:DNA complexes (40).

**Figure 1.**
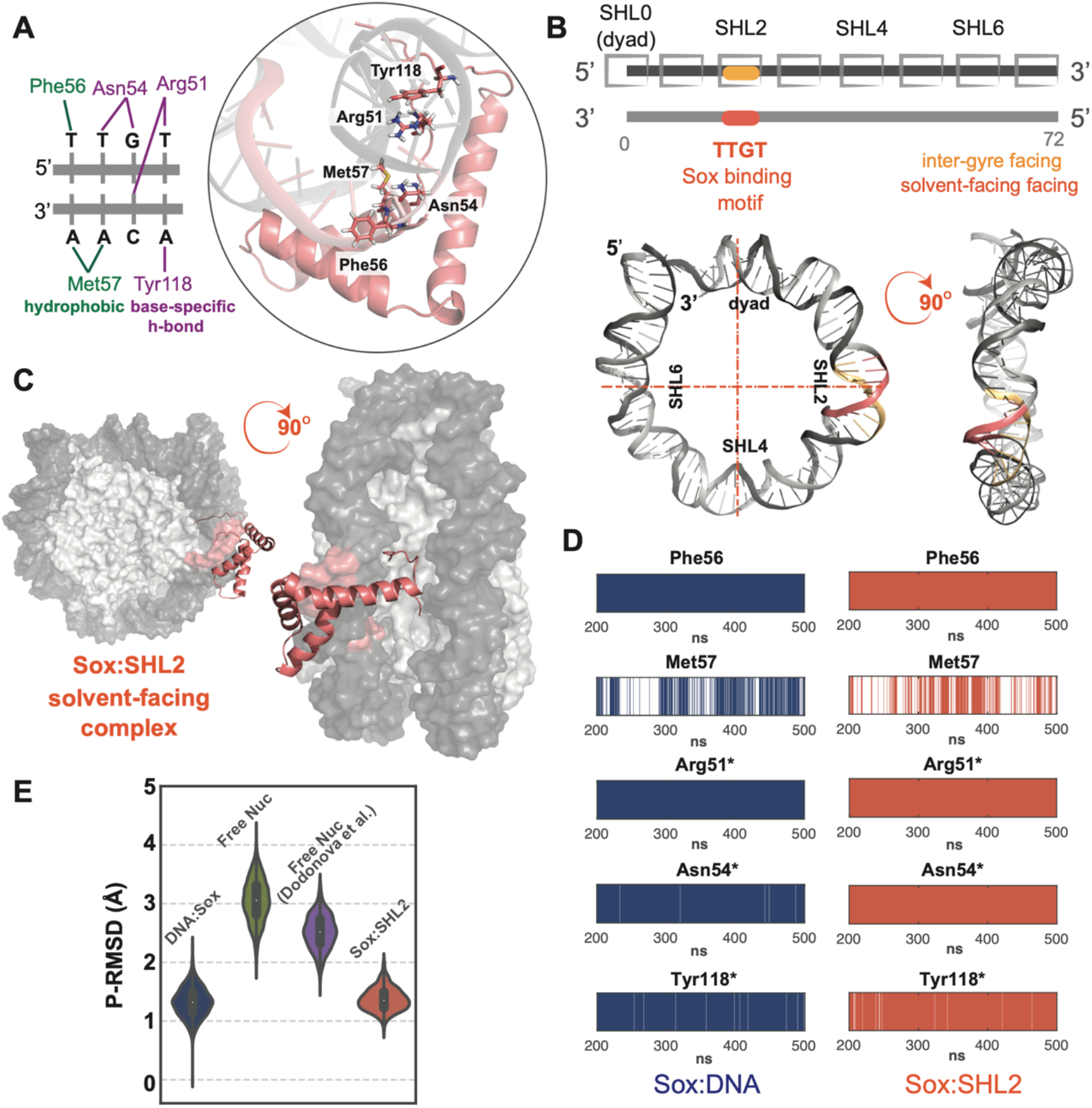
**A. The important Sox:DNA interactions.** The Sox:DNA interactions reported to be critical to Sox:DNA recognition are demonstrated in sticks on Sox11:DNA complex (pdb id: 6t78). The core 5’-TTGT-3’ and its complementary Sox binding motif is specifically recognized by the hydrogen bonds formed by Arg51, Asn54, and Tyr118, and hydrophobic interactions by Phe56 and Met57. Hydrophobic interactions and base-specific hydrogen bonds are colored in green and purple, respectively. **B.** The Sox cognate sequence can be placed either on the solvent-facing or inter-gyre facing strand. The representative coordinates of each positive SHL are highlighted with a box. The corresponding solventfacing (salmon) or inter-gyre-facing (orange) Sox cognate sequence (5’-TTGT-3’) positioning at SHL2 are shown on the nucleosomal DNA structure. **C.** The solvent facing Sox11:SHL2 complex is shown from front and side views, where histones and nucleosomal DNA are depicted as surface and colored in light and dark gray, respectively. Sox is shown as cartoon and colored with salmon. **D.** The Sox11:DNA (dark blue) and Sox11:SHL2 (salmon) interaction profiles of the essential Sox amino acids. Each barcode plot shows the presence of the denoted amino acid interaction with the DNA (within the equilibrated simulation time frame). The prevalent base-specific interactions formed by Arg51, Asn54, and Tyr118 are presented for Sox11:DNA and Sox11:SHL2, respectively and highlighted with an asterisk. **E.** The MD-driven phosphate root mean square deviations (P-RMSDs) from the native Sox-bound-DNA conformation (pdb id 6t78). The P-RMSD distributions of the Sox cognate sequence, derived from Sox11:DNA (dark blue, N=1200), free nucleosomal SHL2 601 DNA (green, N=3000), free nucleosomal SHL2 natural DNA (purple, N=1200), and Sox11:SHL2 model (salmon, N=1200) simulations. The persistence rate of each interaction is provided in Supplementary Table 1.

In 2020, Dodonova *et al.* unveiled the nucleosome recognition mode of Sox2 and Sox11 at the atomistic detail (43). In their complex, Sox cognate sequence was inserted into the nucleosomal DNA at SHL2, where the cognate 5’-TTGT-3’ faces the histone octamer (Figure 1B). At SHL2, Sox2:NCP and Sox11:NCP complexes came out structurally identical (pdb ids: 6t7a, 6t7b). In both complexes, the binding of Sox is accompanied by a strong local distortion of the nucleosomal DNA, involving a 7Å widening of the minor groove. The tight interactions between Sox and nucleosomal DNA resulted in pulling of DNA away from the histone octamer by ~4Å. These DNA perturbations are reminiscent of the ones induced by Sox binding to free DNA, underscoring that Sox:DNA interactions are conserved at SHL2 (44–49). As an intriguing observation, in both structures, 25 bp from the end of the nucleosomal DNA are detached from the histone octamer due to steric clashes formed between Sox and the adjacent gyre (at SHL-5, SHL-6, and SHL-7 positions) (Supplementary Figure 1A-B). At higher Sox concentrations, a second Sox was able to specifically bind to NCP, but this time at SHL-2. This binding resulted in freeing of an additional 25 DNA bp from the other end (at SHL5, SHL6, and SHL7). The binding of Sox appears to reposition the H4 histone tail as well. In this way, Sox could alter the canonical nucleosome-nucleosome contacts to open the chromatin fiber locally (43).

These exciting observations unveiled the fundamental principles of nucleosomal DNA recognition by Sox PTF. However, to completely grasp the universal binding rules of Sox, two further important questions should be answered: (i) Can Sox stably bind to SHL2, even if its cognate sequence is located on the solvent facing strand (the complementary strand to the one used by Dodonova *et. al.);* (ii) Is Sox recognition exclusive to SHL2 or can other SHLs accommodate Sox binding. The exploration of these points requires a detailed investigation of nucleosomal DNA dynamics at different resolutions. For this, we combined *in silico* (integrative modeling and molecular dynamics simulations) and experimental (•OH and UV laser DNA footprinting) techniques that can report on the impact of DNA conformational changes on protein:DNA recognition. Through our *in silico* approaches, we discovered that Sox binds to SHL2 without imposing any major DNA rearrangements when its cognate sequence faces the solvent. We also uncovered that the SHL-specific histone:DNA interactions dictate native Sox recognition, which is only realized at SHL2. These observations could all be validated by •OH and UV laser DNA footprinting techniques, portraying a unified PTF:nuclesome recognition mechanism, guided by the inherent biophysics of the nucleosomes.

## RESULTS

### Molecular dynamics simulations of Sox:DNA complex reveals the essential Sox amino acids for base and shape DNA reading

The earliest Sox:DNA complex dates back to 1995 (50). Since then, a number of other Sox:DNA complexes were characterized. These complexes showed that Sox binds to the minor groove, while inducing a dramatic deformation at its recognition sequence 5’-TTGT-3’ (Figure 1A). In all these complexes, the regions inducing these changes were defined as the hydrophobic FM wedge (Phe56, Met57) and the polar Asn54 (numbering follows the one in Sox11 of 6t78 pdb (43)) (44–46, 48, 49, 51, 52). Among these amino acids, Phe56 triggers the minor groove opening, while the 60°-70° bending is maintained by the base-specific Asn54:DNA interactions with the central 5’-TG-3’ motif (Figure 1A). Next to these shape-preserving essential contacts, the core 5’-TTGT-3’ Sox sequence is further read by Arg51, Asn54, and Tyr118. These three residues form base specific hydrogen bonds with the core and complementary binding motifs, as presented in Figure 1A. To understand the specificity and persistence of these interactions, we performed (two replicas of 500 ns long) molecular dynamics (MD) simulations of Sox11:(free)DNA complex (pdb id: 6t78). As a result, we observed that Arg51, Asn54, and Tyr118 interact with 5’-TTGT-3’ during the whole simulation time in a base-specific manner (Figure 1D-blue profiles, Supplementary Table 1). Furthermore, we saw that from the FM wedge, Phe56 is the one making persistent hydrophobic interactions, while Met57 contacts DNA only for the 50% of the simulation time (Figure 1D-blue profiles, Supplementary Table 1). So, Phe56, Arg51, Asn54, and Tyr118 are indispensable to specific Sox11:DNA recognition. We also observed that the Sox-bound-DNA backbone conformation fluctuates around its initially crystallized state within 0.7-2.2 Å P-RMSD (Phosphorus Root Mean Square Deviation (Methods, Figure 1E – blue distribution)), defining the DNA thermal fluctuation range permitted by Sox binding. We expect that any native Sox:nucleosome complex will reflect the same base-specific contact (base reading) and DNA thermal fluctuation (shape reading) profile. Expanding on this, we explored the dynamic behavior of Sox11-bound nucleosomes, initially at SHL2, and then at dyad and SHL4 positions.

### Only at SHL2 Sox can exert its fingerprint base and shape DNA reading

To probe the universal nucleosomal recognition mechanism of Sox, we structurally modeled a new nucleosome upon inserting the core 5’-TTGT-3’ Sox binding sequence at dyad, SHL2, and SHL4 sites of the 601 nucleosome (pdb id:3lz0, (77)) concurrently. Here, we placed an SHL gap between the Sox binding sequences to ensure that these newly incorporated sites will be distant enough for not to *“feel”* each other. At each mutated SHL site, the core Sox binding sequence was inserted on the solvent-facing nucleosomal DNA strand, i.e., the complementary strand used by Dodonova *et al.* for Sox binding (pdb id: 6t79 (43), Figure 1B, Supplementary Figure 1 A-B). Our motivation in constructing a derivate of the stable 601 nucleosomal DNA sequence was, first, to have a well behaving system for the experimental validation, second, to focus only on the conformational freedom injected to the system upon incorporating Sox binding sites. To explore the thermal fluctuations of the mutated 601 nucleosome beyond the available nucleosomal DNA conformational space (Supplementary Figure 1C-D), we performed two 1 μs long MD simulations. As a result, we observed that irrespective of the nucleosomal positioning, Sox binding sites rarely open wide enough to match the 0.7-2.2 Å P-RMSD fingerprint DNA deformation range (Figure 1E–blue profile, Supplementary Figure F). To serve as a reference, we simulated the natural 6t79 nucleosome too (two 500 ns long MD simulations). At SHL2, this natural nucleosome exerts similar thermal DNA fluctuations as mutated 601 (Figure 1E – green and purple distributions, Supplementary Figure 2B), underscoring the validity of using our mutant nucleosome for further modeling.

Since we know that Sox can recognize its binding sequence at SHL2, from our mutated 601 nucleosome simulations, we isolated the nucleosome state fitting best to Sox-bound-DNA at the SHL2 site (with 1.9 Å P-RMSD, Supplementary Figure 1E). By taking this best fitting SHL2-nucleosome and the bound conformation of Sox11 (from 6t78), we imposed the known Sox11:DNA interactions in HADDOCK and obtained an initial Sox:SHL2 model (78, 79) (Figure 1C, Methods). This model was subjected to two rounds of 500 ns long MD simulations, to allow enough time for Sox11 to find its native binding pose. As a result, we observed that Sox11:SHL2 complex behaves stably during the last 300 ns of the simulations (Supplementary Figure 2A), which we took as a basis for the rest of our analyses. Tracing the P-RMSD profile of Sox11:SHL2 complex and comparing it to the 601-SHL2 P-RMSD profile revealed something very striking: Sox binding is indispensable to induce the extreme minor groove geometry (Figure 1E). So, only when Sox11 is located at its binding site, the P-RMSD distribution of Sox-bound-DNA profiles could be replicated. When we concentrated on the specific interaction profiles of Sox, we observed that at SHL2, the critical amino acids of Sox behave exactly the same as in the case of Sox:DNA complex (Figure 1D). This implies that SHL2 acts transparently to Sox binding, also when Sox cognate motif faces the solvent. This observation tells us that the persistent Sox:DNA interaction network directed by Phe56, Arg51, Asn54, and Tyr118 is required to induce the extreme minor groove deformation also on the nucleosome. As obtaining the Sox:DNA observations on Sox11:SHL2 model required incorporation of integrative structural modeling, as well as running several cycles of MD simulations, we named this protocol after Dynamic Integrative Modeling (DIM, Methods) protocol.

To reveal the impact of histone:DNA interactions on the position-dependent behavior of nucleosomal DNA, we modeled Sox11 binding also at dyad and SHL4 (Figure 2A) following the same DIM protocol, as described above. In the case of Sox:dyad, the P-RMSD profile unveiled that Sox11:dyad almost never exerts the Sox-bound-DNA conformation (Figure 2B). Also, in the case of Sox11:dyad, the specific Asn54 interactions are lost, and the specific Tyr118 interactions are reduced by 50%, while the hydrophobic Met57 interactions are increased by 20% (Figure 2C - left). Interestingly, Sox11:SHL4 complex reflects Sox-bound-DNA conformations during the 50% of the simulation time, which lays out SHL4 as a less favorable binding site compared to SHL2. Additionally, at SHL4, the hydrophobic Met57:DNA interactions are further increased by 40% (Figure 2D). However, in this case, the base-specific Asn54 and Tyr118 hydrogen bonds are almost lost. These findings indicate that the more specific Sox:DNA interactions are lost, the more hydrophobic non-specific interactions prevail. They also imply that SHL4 permits solely shape reading and dyad allows no reading mechanism at all (Supplementary Figure 2B). The denoted interaction profiles were also consistently observed during our replica simulations (Supplementary Figure 2C).

**Figure 2.**
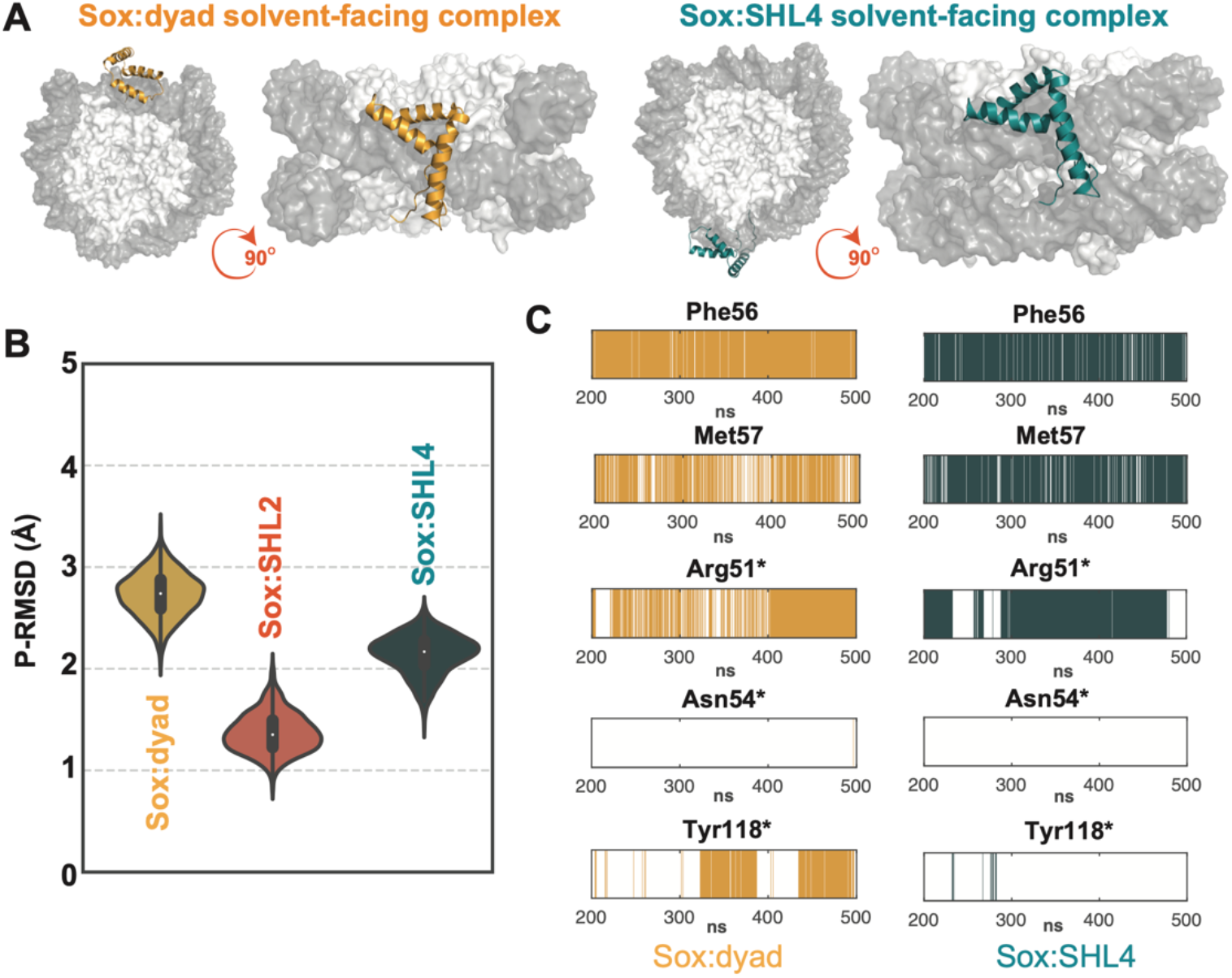
**A.** The solvent facing Sox11:dyad (orange) and Sox11:SHL4 (green) complex models. The models are shown from front and side views, where histones and nucleosomal DNA are represented in surface and colored in light and dark gray, respectively. Sox is presented in orange and green cartoon for Sox11:dyad and Sox11:SHL4 binding, respectively. **B.** The MD-driven P-RMSDs of each Sox11:SHL complex compared to the Sox-bound-DNA conformation. P-RMSD distributions of the Sox cognate sequence, derived from Sox11:dyad (orange, N=1200), Sox11:SHL2 (salmon, N=1200), Sox11:SHL4 (dark green, N=1200) complex simulations. The Sox11:SHL2 values are a replicate of Figure 1E and placed here to serve as a basis for a fair comparison. **C.** The Sox11:DNA interaction profiles of the essential Sox amino acids at dyad and SHL4 (the data comes from a population where N=600). Each barcode plot shows the presence of the denoted amino acid interaction with DNA (within the equilibrated simulation time frame). The prevalent the base-specific interactions formed by Arg51, Asn54, and Tyr118 are highlighted with asterisk. The persistence rate of each interaction is given in Supplementary Table 1.

### Solvent-facing Sox binding is confirmed by the •OH footprinting experiments

To confirm the solvent-facing Sox binding at the dyad, SHL2, and SHL4, we used hydroxyl radical (•OH) footprinting. The •OH footprinting is a versatile technique to analyze the binding of Sox to NCP, as •OH radicals attack DNA via the minor groove (53–55). As Sox binds to the DNA minor groove, a footprint should be visible at each relevant SHL site. Expanding on this, we performed •OH footprinting on our mutated 601 sequence, in isolation and when wrapped around the histones (Methods, Figure 3A and Supplementary Figure 3). In these experiments, instead of Sox11, we used the Sox6 to trace the Sox binding, as it stably behaved during our experiments. Even though Sox6 does not have any experimentally determined structure, we permitted this as Sox binding to DNA is strictly conserved across the Sox family. This is also reflected by the identity and similarity percentages of the HMG domains shared between Sox11 and Sox6 (59% and 86%, respectively). During our experiments, both naked DNA and nucleosomes were allowed to interact with the increasing amount of Sox (the amount of Sox used for the analysis of Sox binding to the NCP was ~ 3-fold higher than used in the naked DNA experiments). Afterwards, the samples were used for •OH radical footprinting. As seen in Figure 3A, upon increasing the amount of Sox in the reaction mixture, a very clear protection of all three Sox recognition sequences is observed on the naked DNA (Figure 3A). Protection is also observed at Sox binding sequences in histone-wrapped mutated 601 DNA (Figure 3A, insets). Albeit the protection at SHL2 and SHL4 is not so well visible. Here, the protected sites coincide with the maximal cleavage of the DNA in the nucleosome. Of note, a very specific footprinting of Sox is detected at the nucleosome dyad, where the four middle DNA bands exhibit a strong decrease in the intensity compared to the flanking bands (see the magnified recognition sequence footprint as well as the scans at the right inset of Figure 3A). This Sox-protected •OH cleavage pattern is very similar to the one of globular linker histone H1 domain in the H1-bound nucleosome ((2, 4, 56), see also Supplementary Figure 4A-B), which implies that the global 3D organization of Sox-dyad complex is analogous to the complex formed between the globular domain of H1 and the nucleosome dyad (2, 4, 56).

**Figure 3.**
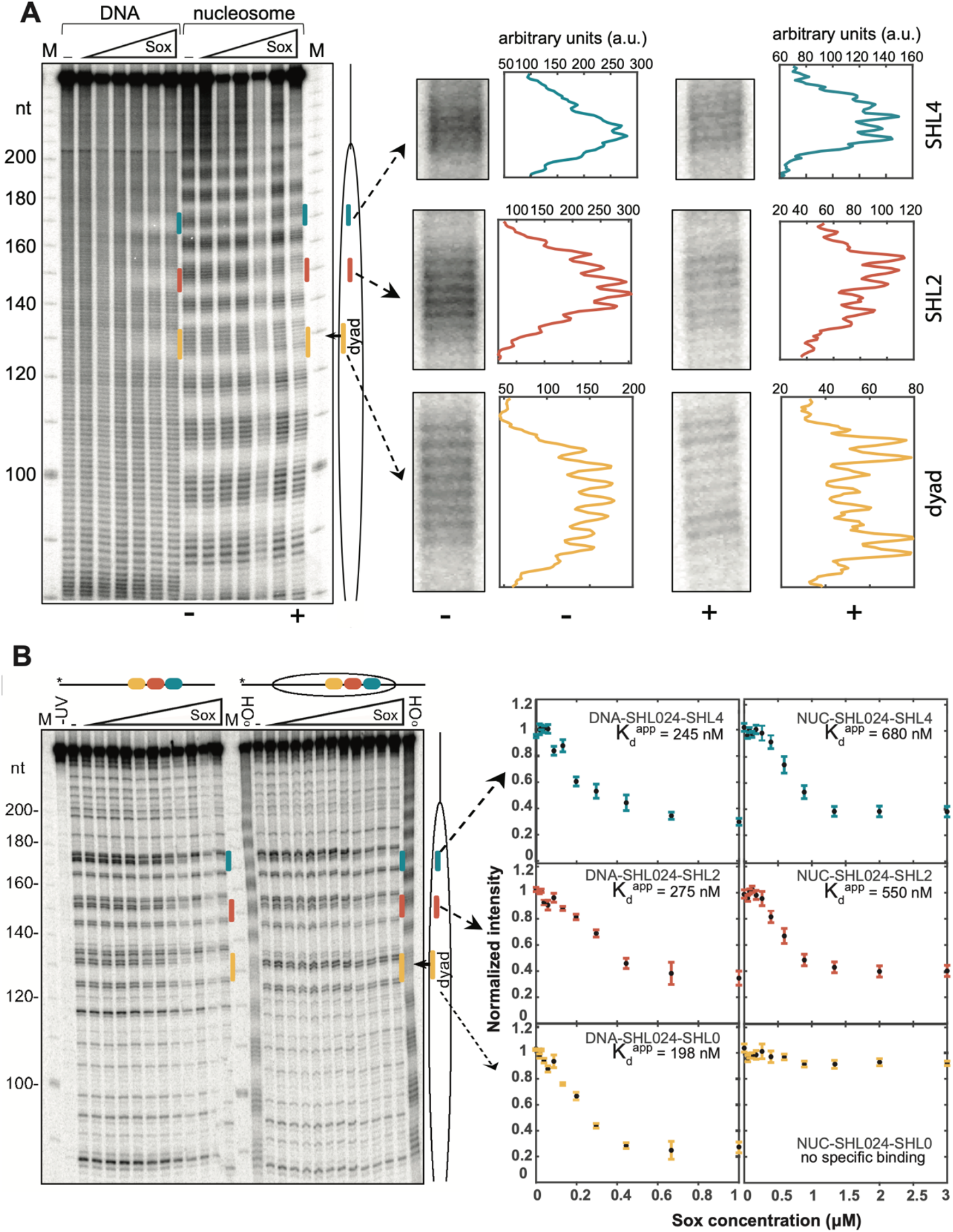
**A.** Three Sox recognition sites that are accessible to Sox binding within the nucleosome. ·OH radical footprinting patterns of Sox-DNA and Sox:nucleosome complexes, bearing the three recognition sites of Sox (601-SHL024). After ·OH treatment of the complexes, the cleaved DNA fragments were purified and separated on 8% sequencing gel. They were then visualized by autoradiography. (−, +) sign means that the two gel lanes are zoomed (−) and (+) Sox6 protein, i.e. the first and the last lane in the nucleosome part of the gel (left). Right, higher magnification of the indicated Sox binding sites. ·OH cleavage patterns of in un-bound (−) and Sox6-bound particle (+), containing the three Sox recognitions sites. Vertical red lines correspond to Sox binding sites; oval to the schematics of the nucleosome, where the dyad is indicated with an arrow. **B.** Left Panel: UV laser footprinting patterns of the Sox6-DNA and Sox6-nucleosome complexes bearing a Sox recognition sequence at 601-SHL024. The complex was irradiated with a single 5 nanoseconds UV laser 266 nm pulse (Epulse,0.1 J/cm^2^) and DNA was purified from the samples. After treatment of the purified DNA with Fpg glycosylase, the cleaved DNA fragments were separated on 8% sequencing gel and visualized by autoradiography. Red vertical lines and red squares mark the Sox binding sites, M marks the molecular mass, oval represent the schematics of the nucleosome, and the dyad is indicated with an arrow. -UV refers to the control, non-UV irradiated, and Fpg treated sample. Right panel: (left) Sox6 concentration dependencies of the footprinting intensities, representing the normalized cleavage band intensity of the GG run within the binding site. (right) The normalized intensity profiles based on our gel quantifications were plotted as a function of Sox6 concentration.

### UV laser footptinting validates that DNA shape reading is realized at SHL2 and SHL4, but not at the dyad

To analyze the local DNA changes occurring at the minor groove of nucleosomal DNA, we performed UV laser footptinting of Sox-bound mutated 601 DNA in free and histone-wrapped states. UV laser footprinting measures the UV laser-induced alterations in the nucleotide photoreactivity (57), which could affect the spectrum and the amount of the DNA lesions. Since the binding of Sox to DNA alters the local structure of Sox recognition sequence, it should lead to changes in the spectrum of lesions. Such lesions are extremely sensitive to local DNA structure and can easily be mapped by alkali or enzymatic DNA strand cleavage, followed by electrophoresis under denaturing conditions at the single nucleotide resolution (58). Since a single nano- or picosecond laser pulse is used for irradiation, the generation of the lesions is achieved in an interval of time, which is shorter than the conformational transitions of the protein:DNA complex (59). So, the laser footprinting is taking a snapshot of the complex structure, while recording the Sox-induced structural signature (59). We used this method successfully in the past for mapping productive protein:DNA interactions (17, 60, 61). Followingly, we mapped UV laser specific biphotonic 8-OxoG lesions by using Fpg glycosylase (formamidopyrimidine [fapy]-DNA glycosylase). These lesions are observed on a GG sequence, located right after the 5’-ACAA-3’, complementary to Sox cognate 5’-TTGT-3’ (Supplementary Figure 5A-B and Methods). As expected, with the increase in Sox concentration, the disappearance of 8-oxoG bands is traced in the mutated 601 (601-SHL024) DNA constructs, indicating a deformation at the Sox cognate sequence (Figure 3B, see also Supplementary Figures 3A-B and 4C). In the case of 601-SHL024 nucleosome, the same behavior was observed at SHL2 and SHL4, while at the dyad no footprint was spotted (Figure 3B). Also, apparent dissociation constant (K_d_^app^) values of specific Sox binding to its target sequence (Table 1) were evaluated by Sox titration footprinting (Figure 3B (right) and Supplementary Figure 6) and gel quantification (Supplementary Figure 5C). Interestingly, the measured apparent binding affinities for SHL2 and SHL4 nucleosomes were only 3-4 times lower than in naked DNA. The higher binding affinity of SHL2 compared to SHL4 in nucleosomes, together with the absence of specific binding at the dyad, is fully consistent with the data obtained by our DIM approach. Important to note that this apparent (measured) dissociation constant (K_d_^app^) must not be associated with the true dissociation K_d_. Indeed, the two orders of magnitude higher concentrations (R=50 nM) of the constant component (DNA or nucleosomes) with respect to the true K_d_ <1 nM (62) and the unavoidable presence of several lower-affinity binding sites per ~200 bp DNA fragment (due binding motif degeneracy) precluded the true K_d_ determination under our experimental conditions (see (63)). All in all, our findings revealed that Sox can produce similar local conformational changes, i.e., minor groove widening, at SHL2 and SHL4 as in free DNA, confirming our modeling and simulation results. Identical footprinting profiles were observed when the Sox cognate sequence is incorporated at the dyad, SHL2, and SHL4 separately (Supplementary Figure 5C and 6). We should also note that our computational results for Sox11 were reproduced when we performed our DIM protocol with a homology model of Sox6, which endorses the general applicability of our findings (Methods, Supplementary Figure 7).

**Table 1.** The apparent dissociation constants (K_d_^app^) of Sox6 binding to naked and nucleosomal DNA representing the concentrations of Sox6 at the half maximum signal intensity change, extrapolated by the least square fitting procedure of data in Figure 3B (right) and Supplementary Figure 6. * denotes the absence of binding to the target sequence.

| SHL region | DNA<br>$K_d^{app}$ (nM) | Nuc<br>$K_d^{app}$ (nM) |
| --- | --- | --- |
| SHL0 | 100 | * |
| SHL2 | 136 | 374 |
| SHL4 | 180 | 490 |
| 3x SHL0 | 198 | * |
| 3x SHL2 | 275 | 550 |
| 3x SHL4 | 245 | 680 |

## DISCUSSION

PTFs have the remarkable ability to directly bind to chromatin for stimulating vital cellular processes. Thus, PTF binding should be universal, where the strand-positioning of PTF cognate sequence should not impact its accessibility on the nucleosomes. Earlier, Sox binding to histone-facing cognate sequence was determined by cryo-EM (43). In this EM structure, the solvent-excluded positioning of the Sox recognition motif creates a steric clash between the two helices of Sox and the two DNA gyres (Supplementary Figure 1A). In parallel to this finding, in this work, we studied how Sox PTF binds to a solvent-facing nucleosomal DNA. Accordingly, in our Sox:nucleosome model, we did not observe any major structural changes, as in this mode Sox can comfortable fit in between DNA gyres (Figure 1D, Figure 2A). Strikingly, our model and Dodonova *et. al.’s* Sox:nucleosome structure are of the same accuracy (P-RMSD: 1.4 Å), endorsing the validity of our approach.

In this work, we also showed for the first time that only SHL2 acts transparently to Sox binding. This is not a surprise, as it was already demonstrated that the nucleosome displays stretching at SHL2 with a kink into the minor groove, providing a perfect recognition site for proteins (64). Relatedly, it was recently shown that SHL2 can harbor nonspecific protein binding, as revealed by the TATA-box-binding-protein:SHL2 complex (65). At the SHL2 H3 α3- and H4 α2-helices form very few protein contacts, providing an extra flexibility to this region (66–69). All in all, it is clear that weak histone-DNA interactions at SHL2 makes SHL2 a good recognition site for Sox. These observations were also validated by our UV laser footprinting experiments.

Further structural analysis on our Sox11:SHL complexes showed that only at SHL2, Sox’s globular α-helical HMG domain is far enough for not to directly feel the basic nature of histones (Figure 4A, Supplementary Table 2). This is in consistent with the fact that the most stable Sox11 binding (with the least thermal fluctuations) was observed at the SHL2 site (Figure 4B). At the dyad, which is the least stable binding site, the histones fix Sox-DNA motif in a symmetric fashion, preventing the asymmetric bending of the cognate DNA. Interestingly, this is not observed at SHL4, as the non-symmetric histone-DNA interactions at the SHL4 permits the asymmetric deformation of Sox-DNA. However, in this case, the N-terminal H2A Lys15 interferes with the conserved His75 of Sox11, leading to a local destabilization at the tip of the Sox (Figure 4B). All these outcomes explain how SHL2 could act as a “transparent” hot spot for Sox binding. In the light of these results, we claim that the histone-DNA interactions are tuned at SHL2 to allow both shape and base reading of DNA, whereas at SHL4 only a non-specific shape reading could be achieved (Supplementary Movies 1-3).

**Figure 4.**
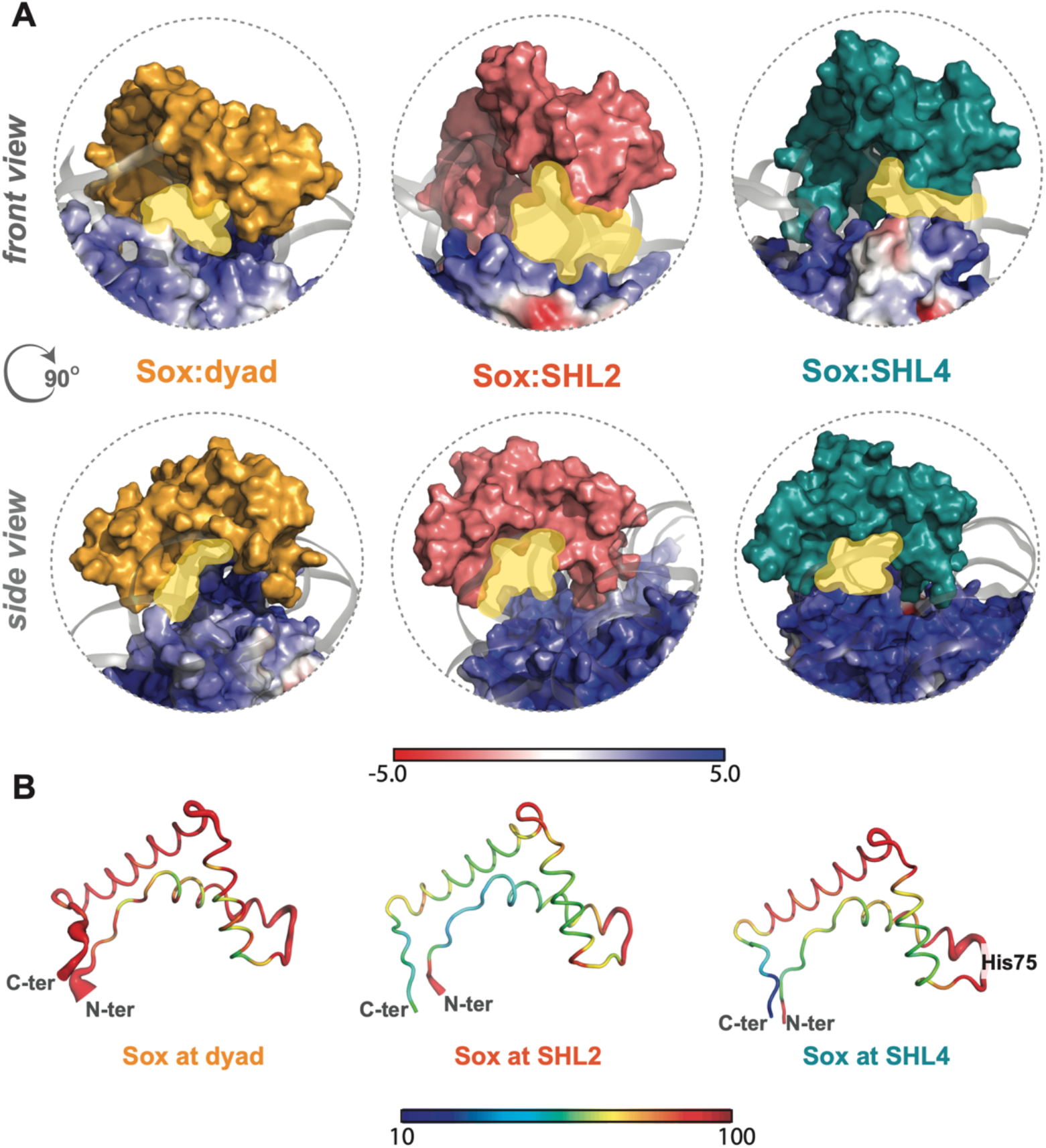
**A.** SHL2 is transparent to Sox binding. Sox is far enough from the basic histones only at the SHL2 binding site. The nucleosomal DNAs are demonstrated as cartoon and colored in dark gray. Histones are shown as surface and colored according to their electrostatic potentials. Here, red represents negatively charged (−5) regions and blue represents positively charged (+5) regions. **B.** Sox binds the most stable at SHL2. The Sox PTFs deduced from the MD simulations of Sox11:dyad (left), Sox11:SHL2 (middle) and Sox11:SHL4 (right) are colored according to their temperature factors. Blue and red colors depict the most rigid and the most flexible regions, respectively. The cartoon thickness linearly scales with the degree of thermal fluctuations (averaged over N=600 conformations). When Sox11 is bound to the SHL2, it has the most stable conformation.

Also, through our experiments, we observed that K_d_^app^ has lower values when both base and shape reading are satisfied, as in SHL2 (Figure 1D-E and Figure 3B). However, more detailed experiments are required to comment on how Sox binding scales among different SHLs. Finally, we disclosed that •OH radical footprinting of Sox at the dyad is very similar to the H1 globular domain bound at the dyad (Supplementary Figure 4A-B). This finding suggests that different HMG proteins could be able to efficiently bind *in vivo* to the dyad, without requiring the presence of the recognition motif. In other words, the peculiar V-shape of the entry-exit nucleosomal DNA (2, 56). Thus, very similar shape and globular H1 domain-like dimensions of Sox HMG-domain could be sufficient for non-specific binding of HMG-domain proteins to the dyad. This could explain why the non-specific and highly abundant HMGB1/B2 chromatin proteins and the linker histone would have a shared structural role in organizing linker DNA in the nucleosome (70).

In summary, our *in silico* and experimental observations in combination reveal that the binding of Sox is strongly nucleosomal-context-dependent, where histone-DNA interactions dictates the binding capacity of Sox, which could be valid for other PTFs as well.

## METHODS

### Dynamic integrative modeling

To explore Sox:nucleosome interactions (both for Sox11 and Sox6), we performed a series of modeling and molecular dynamics simulation cycles, making up our Dynamic Integrative Modeling (DIM) approach. Our DIM protocol is composed of three steps (Supplementary Figure 8): (**Step 1**) Inserting the Sox cognate sequence into 601 Widom DNA sequence (pdb id:3lz0, (77)) at different SHLs, i.e., SHL0 (dyad), SHL2 and SHL4, followed by its molecular dynamics (MD) simulations (free mutated nucleosome). (**Step 2**) Isolating the Sox-binding-compatible nucleosome conformer for each SHL to model Sox-bound nucleosomes. (**Step 3**) Simulating the constructed Sox-bound nucleosome structures. We also simulated the Sox11:(free)DNA complex (pdb id:6t78, (43)) to serve as a reference. The technical details of each step are provided below.

(**Step 1**) The MD simulations of the 601 Widom DNA sequence: We inserted a special sequence, bearing the Sox consensus sequence 5’-GG<u>*ACAA*</u>TGGAGG-3’ at dyad, SHL2 and SHL4 by using 3DNA (74). The mutated sequence corresponds to (−4^th^)-7^th^, 16^th^-27^th^ and 37^th^-48^th^ nucleotides in the forward DNA chain, leading to:

5’-ATCAGAATCCCGGTGCCGAGGCCGCTCAATTGGTCGTAGACAGCTCTAGCACCGCTTAAACGCA *CGTAG<u>GACAATGGAGG</u>CGCGTTTT<u>GGACAATGGAGG</u>CATTACTCC<u>GGACAATGGAGG</u>CACGTG* TCAGATATATACATCGA-3’

The mutant nucleosome is named as 601-SHL024, which was subjected to MD simulations, totaling 2 μs long (2 x 1 μs). The conformations with the lowest P-RMSD values were selected to model the Sox bound nucleosomes.

(**Steps 2&3**) The modeling and simulation of the Sox:NCP complexes: The free nucleosome conformers, reflecting the lowest P-RMSD at the dyad, at the SHL2, and at the SHL4 were isolated. Sox11 isolated from the Sox11:DNA complex (pdb id: 6t78, (43)) was located at three locations by template-based modelling. The initial crude Sox:nucleosome complexes were refined within HADDOCK 2.2 (78, 79), while imposing the critical Sox:DNA interactions as restraints (80). Here, we imposed the crystal Sox11:DNA distances, measured between 56^th^, 57^th^ (FM) residues of Sox11 and 0^th^, 20^th^, and 41^st^ nucleotides (forward strand), respectively. This restraint ensures the reproduction of the hydrophobic wedge interactions. The best HADDOCK models were then subjected to two parallel MD simulations, totaling 1 μs. At the end, the conformations with the lowest P-RMSDs were isolated and kept as final Sox11 bound nucleosome structures (for Sox:dyad, Sox:SHL2, and Sox:SHL4). Finally, to serve as a reference, Sox11:DNA complex (pdb id: 6t78) was subjected to two independent MD simulations, totaling 1 μs. The same procedure was carried out with the homology model of Sox6, where for each complex one 0.5 μs MD run was set.

### Molecular dynamics simulations and analysis protocols

All the MD simulations were performed by using GROMACS simulation packages (Gromacs 5.1.4, Gromacs 2019, and Gromacs 2020.4) (71) under the effect of Amber Parmbsc1 force field (72). We used TIP3P as the water model. The NaCl concentration was kept as 0.15 M. A dodecahedron simulation box was used, while having a minimum distance of 12 Å between the biological molecule and the edges of simulation box. The temperature was kept at 310K throughout the simulation. We carried out two replica simulations of the free 601 nucleosome (pdb id: 3lz0) (2 x 1 μs), free natural nucleosome (pdb id: 6t79) (2 x 0.5 μs), the Sox11:DNA complex (pdb id: 6t78) (2 x 0.5 μs), and the Sox11-bound nucleosome models, i.e., Sox11:dyad (2 x 0.5 μs), Sox11:SHL2 (2 x 0.5 μs), and Sox11:SHL4 (2 x 0.5 μs). We also carried out a single simulation for the Sox6-bound nucleosomes i.e., Sox6:dyad (1 x 0.5 μs), Sox6:SHL2 (1 x 0.5 μs), and Sox6:SHL4 (1 x 0.5 μs). Here, Sox6 was a homology model, as there was no experimentally determined structure available. The homology model of Sox6 was generated with Modeller (73).

Before running the simulations, complexes were minimized by using the steepest descent algorithm in the vacuum. Then, they are solvated in TIP3P water and concentration was kept at 0.15 M by adding NaCl to the system (460 Na+, 240 Cl- for free 601 nucleosome; 421 Na+, 201 Cl- for free natural nucleosome, 49 Na+, 38 Cl- for Sox11:DNA; 427 Na+, 218 Cl- for Sox11:dyad; 438 Na+ and 229 Cl- for Sox11:SHL2; and 477 Na+ and 268 Cl- for Sox11:SHL4). The number of ions were added to the topology files accordingly and then the solvated systems were minimized. The systems were relaxed for 20 ps at 310K under the constant volume. To generate replicas, random seeds were changed. Then, another 20 ps molecular dynamics simulations were performed under constant pressure at 1 bar. Finally, position restrains were released gradually from 1000 to 100, 100 to 10 and 10 to 0. The integration time step was set to 2 fs. For the analysis, coordinate files were recorded in every 0.5 ns. The initial 200 ns of all simulations (except for the free 601 nucleosome) were set as the equilibration time of the system and discarded before the analysis stage. The equilibration time for the 601 free nucleosome was set as 350 ns.

At the end of MD simulations, minor groove widening and P-RMSD (Root Mean Square Deviations of the DNA Phosphorus atoms) metrics were calculated over all the conformers generated. Minor groove widths were measured with 3DNA (74). P-RMSD values were computed over the phosphorus atoms of seven nucleotides’, involving the Sox recognition sequence (5’-GACAATG-3’). The reference seven nucleotides correspond to (−3rd)-3rd, 17th-23rd, 38th-44th nucleotide positions at dyad, SHL2 and SHL4, respectively. P-RMSD values of free nucleosome determined by Dodonova et al. measured at SHL2 position. During all these measurement the DNA of the Sox11:DNA complex (pdb id: 6t78, (43)) was taken as a reference. The fitting and P-RMSD computations were performed with Profit (Martin, A.C.R., http://www.bioinf.org.uk/software/profit/).

Molecular interaction profiles of complexes were calculated on each replica simulation by using the Interfacea Python library (https://github.com/JoaoRodrigues/interfacea)(75). This library provides the intra- and inter-interactions of complexes as hydrophobic and ionic interactions and h-bonds in a .txt file for every coordinate file. The output files give the interacting atoms and residue pairs. From these output files, we extracted the interactions between essential Sox amino acids (Arg51, Asn54, Phe56, Met57, Tyr118) and DNA. We isolated the base-specific interactions among all interactions, which are the interactions between the Sox protein and the respective DNA bases. The barcode plots were generated as in the following. If there is an interaction between the selected Sox amino acid and the base of DNA, there is a line drawn in the barcode graph and otherwise the position for a given time is left blank. These graphs were plotted in MATLAB R2020B (76).

### Sox6 HMG domain cloning and purification

The HMG domain of human Sox6 (618 - 697 amino acids) gene was cloned in pET28b vector in between NdeI and XhoI restriction sites. The N terminal His-tagged Sox6 HMG domain was produced in *Escherichia coli* BL21 (DE3) pLYsS cells. Briefly 200 ng of plasmid was used to transform into *E. coli* cells, after transformation bacteria were platted on LB agar plates supplemented with kanamycin & chloramphenicol, and incubated overnight at 37°C. Single colonies were added to 3 ml of LB medium (kanamycin+ chloramphenicol) for 12 to 16 hours at 37°C under shaking at 200 rpm speed. 1ml of amplified bacteria was added to 300 ml LB (kanamycin+ chloramphenicol) and left overnight at 37°C and 200 rpm. For each liter of LB (kanamycin+ chloramphenicol) needed, 10 ml of transformed bacteria were added. After 3 hours of incubation, OD at 600 nm was measured. If the OD600 was comprised between 0.5 and 0.6, bacteria were induced with 0.2 mM IPTG (isopropyl-beta-d-thiogalactopyranoside) at 37°C for 3-4 hours at 200 rpm. After induction bacteria were pelleted at 5000g for 20 min at 4°C. The recombinant human Sox6 HMG domain was purified from supernatant of bacterial lysate by using NiNTA resin (Complete His-Tag purification Resin, Roche) and followed by SP sepharose column chromatography (GE Healthcare). Purity of purified HMG doman of Sox6 protein was analyzed by using 18% SDS-PAGE and stained with coommassie blue.

### Core histone purification

Human histones H2A, H2B and H3 were sub-cloned in pHCE vector system and human histone H4 sub-cloned in pET15b vector system. Histones H2A, H2B and H3 were produced in *Escherichia coli* BL21(DE3) cells and human H4 was produced in *E. coli* JM109(DE3) cells. Core histones were produced as N-terminal His-tagged proteins in *E. coli* cells in the absence of T7 RNA polymerase by omitting the addition of isopropyl-P-D-thiogalactopyranoside, which induces the T7 RNA polymerase production in BL21(DE3) and JM109(DE3) cells. Briefly 200 ng of plasmid (for each histone) were used to transform into respective *E. coli* strains. 10 colonies were inoculated into 2L LB broth (ampicillin final concentration 50μg/ml) in 5L flask and left overnight at 37°C and 200 rpm. Each liter of bacteria was pelleted at 5000g for 20 min at 4°C. The cells producing recombinant histones were collected and disrupted by sonication in 50 ml of buffer A (50 mM Tris-HC1 (pH 8.0), 500 mM NaC1, 1 mM PMSF, and 5% glycerol). After centrifugation (27,216 x g; 20 min; 4°C), the pellet containing His-tagged histones as insoluble forms was resuspended in 50 ml of buffer A containing 7 M guanidine hydrochloride. After centrifugation (27,216 x g; 20 min; 4°C), the supernatants containing the His-tagged histones were combined with NiNTA resin (Complete His-Tag purification Resin, Roche) (1mL of Ni-NTA per 1L of bacteria) and were mixed by rotation for 1 hr at 4 °C. The agarose beads were packed into an Econo-column (Bio-Rad) and were then washed with 100 ml of buffer B (50 mM Tris-HC1 (pH 8.0), 500 mM NaC1, 6 M urea, 5 mM imidazole, and 5% glycerol). The His-tagged histones were eluted by a 100 ml linear gradient of imidazole from 5 to 500 mM in buffer B, and the samples were dialyzed against buffer C (5 mM Tris-HC1 (pH 7.5) and 2 mM 2-mercaptoethanol).

The N-terminal 6x His tags were removed from the histones by thrombin protease (GE Healthcare) treatments using 1 unit per 1 mg of protein for 3-5h at 4°C. The removal of the His tags was confirmed by SDS-16% polyacrylamide gel electrophoresis (PAGE); the recombinant histones without the His tag migrated faster than the His-tagged histones. After uncoupling of the His tag, each histone was subjected to Resource S column chromatography (GE Healthcare). The column was washed with buffer D (20 mM sodium acetate (pH 5.2), 200 mM NaC1, 5 mM 2-mercaptoethanol, 1 mM EDTA, and 6 M urea), and each histone was eluted by a linear gradient of NaC1 from 200 to 900 mM in buffer D. The fractions containing the pure histone were mixed and stored in -80°C.

### Histone tetramers and dimers preparation

To prepare tetramers and dimers human H3 & H4 and human H2A & H2B were mixed in an equimolar ratio, and dialyzed overnight in HFB buffer (2M NaCl, 10 mM Tris pH7.4, 1 mM EDTA pH 8 and 10 mM β-mercaptoethanol). After dialysis the supernatant containing folded tetramers and dimers were subjected to Superose 6 prep grade XK 16/70 size exclusion column (GE Healthcare) purification using HFB buffer. The major fractions containing purified tetramers and dimers were mixed. For long time storage, tetramers and dimers were mixed with NaCl saturated glycerol to achieve the final glycerol concentration around 15-20% and stored at -20°C.

### Preparation of DNA fragments

The 255 bp of 601 DNA constructs containing Sox6 consensus motif 5’-GGACAATGGAGG-3’ positioned in different places were produced by chemical synthesis method and cloned into standard vector pEX-A by Eurofins Genomics, Germany. The position of Sox6 binding sites in the 601 constructs were mentioned below,

Sox-SHL0 (dyad): (Sox6 binding motif located at 66bp away from the end of nucleosomal DNA)

GCATGATTCTTAAGACCGAGTTCATCCCTTATGTGATGGACCCTATACGCGGCCGCCATCAGAAT CCCGGTGCCGAGGCCGCTCAATTGGTCGTAGACAGCTCTAGCACCGCTTAAACGCACGTA<u>GGACAATGGAGG</u>CGCGTTTTAACCGCCAAGGGGATTACTCCCTAGTCTCCAGGCACGTGTCAGATAT ATACATCGATGTGCATGTATTGAACAGCGACCTTGCCGGTGCCAGTCGGATAGAATTCCGGAC

Sox6-SHL2: (Sox6 binding motif located at 46bp away from the end of nucleosomal DNA)

GCATGATTCTTAAGACCGAGTTCATCCCTTATGTGATGGACCCTATACGCGGCCGCCATCAGAATCCCGGTGCCGAGGCCGCTCAATTGGTCGTAGACAGCTCTAGCACCGCTTAAACGCACGTACGC GCTGTCCCCCGCGTTTT<u>GGACAATGGAGG</u>CATTACTCCCTAGTCTCCAGGCACGTGTCAGATAT ATACATCGATGTGCATGTATTGAACAGCGACCTTGCCGGTGCCAGTCGGATAGAATTCCGGAC

Sox6-SHL4: (Sox6 binding motif located at 25bp away from the end of nucleosomal DNA)

GCATGATTCTTAAGACCGAGTTCATCCCTTATGTGATGGACCCTATACGCGGCCGCCATCAGAATCCCGGTGCCGAGGCCGCTCAATTGGTCGTAGACAGCTCTAGCACCGCTTAAACGCACGTACGC GCTGTCCCCCGCGTTTTAACCGCCAAGGGGATTACTCC<u>GGACAATGGAGG</u>CACGTGTCAGATAT ATACATCGATGTGCATGTATTGAACAGCGACCTTGCCGGTGCCAGTCGGATAGAATTCCGGAC

Sox6-SHL024: (Sox6 binding motifs located at 66bp, 46bp and 25bp away from the end of nucleosomal DNA)

GCATGATTCTTAAGACCGAGTTCATCCCTTATGTGATGGACCCTATACGCGGCCGCCATCAGAAT CCCGGTGCCGAGGCCGCTCAATTGGTCGTAGACAGCTCTAGCACCGCTTAAACGCACGTA<u>GGACAATGGAGG</u>CGCGTTTT<u>GGACAATGGAGG</u>CATTACTCC<u>GGACAATGGAGG</u>CACGTGTCAGATAT ATACATCGATGTGCATGTATTGAACAGCGACCTTGCCGGTGCCAGTCGGATAGAATTCCGGAC

All Sox6 binding motif harboring 601 constructs were amplified using ^32^P end labeled primers. The labeled DNA substrates were purified on 5% native acryl amide gel prior to use for nucleosome reconstitutions.

### Nucleosome Reconstitution

Nucleosome reconstitution was performed by the salt dialysis procedure. Approximately, 250ng of 32P-labeled DNA probe containing the Sox6 binding site and 4.5 μg of chicken erythrocyte DNA (150– 200 bp) as carrier were mixed with human histones-tetramers and dimers approximately in 1: 0.5: 0.5 ratio in HFB buffer (2M NaCl, 10 mM Tris pH7.4, 1 mM EDTA pH 8 and 10 mM β-mercaptoethanol) respectively. The mixtures were transferred into dialysis tubing and the reconstitution was done by dialysis against a slowly decreasing salt buffer. The NaCl concentration starts at 2 M and decreases slowly up to 500 mM NaCl. Indeed, with the help of a peristaltic pump, low salt buffer is added to the high salt buffer beaker at the rate of 1.5 ml/min for 18 h. Once finished, the dialysis bags were transferred to a 300 mM NaCl buffer and left for buffer exchange for 2 h, which was followed by a final dialysis in 10 mM NaCl buffer overnight. All NaCl buffers for reconstitution include 10 mM Tris pH 7.4, 0.25 mM EDTA, 10 mM β-mercaptoethanol and the desired amounts of NaCl.

### Sox6 HMG domain binding reaction

The binding reaction of Sox6 HMG domain on DNA or nucleosomes was carried out at 37°C. Typically, Sox6 HMG domain was mixed with DNA or nucleosome (50nM) in a 20 μl reaction containing 1X binding buffer (10 mM Tris, pH 7.4, 75 mM NaCl, 1 mM EDTA, 1 mM DTT, 100 mg/ml BSA, 0.01% NP40 and 5% glycerol). The naked DNA was supplemented with carrier nucleosomes to a final concentration equal to those of labeled nucleosomes. The maximal Sox6 concentration was 1000 nM and 3000 nM in naked DNA and nucleosome samples respectively and the dilution step was 1.5. An aliquot of this reaction mix was used to check the formation of the Sox6 HMG domain:DNA or Sox6 HMG domain:nucleosome complex by 5% native PAGE at room temperature in 0.3X Tris-borate-EDTA (TBE) buffer. The remaining aliquots were probed by UV laser footprinting.

### Hydroxyl radical footprinting

Hydroxyl radical footprinting was carried out to check the strong nucleosome positioning ability of Sox6 binding site incorporated 255bp of 601 constructs. The reaction was carried out in 15 μl final reaction mixture in quencher free buffer placed at the bottom of an eppendorf tube. The hydroxyl radicals were generated by mixing 2.5 μl each of 2 mM FeAmSO4/4 mM EDTA, 0.1 M ascorbate, and 0.12%H2O2 together in a drop on the side of the reaction tube before mixing rapidly with the reaction solution. The reaction was terminated after 2 minutes by addition of 100 μL stop solution (0.1% SDS, 25 mM EDTA, 1% glycerol, and 100 mM Tris, pH 7.4), and the DNA was purified by phenol/chloroform extraction and ethanol/glycogen precipitation. The DNA was resuspended in formamide loading buffer, heated for 3 minutes at 80°C and ran along with UV laser samples on 8% denaturing gel in 1X TBE buffer. The gels were dried and exposed overnight on a phosphor imager screen. The gels were scanned on phosphor imager and analyzed by Multi-Gauge (Fuji) software.

### UV laser footprinting

The UV laser specific biphotonic lesions 8-oxoG were mapped by Fpg glycosylase which is generated in the Sox6 cognate binding sequence upon UV laser irradiation. The samples were exposed to a single high intensity UV laser pulse (Epulse ~0.1 J/cm2) as described in previous studies (Angelov et al., 2004; Lone et al., 2013). The DNA was then purified by phenol-chloroform and ethanol/glycogen precipitated. The purified DNA was resuspended in resuspension buffer (10 mM Tris, pH 7.4, 30 mM NaCl, 1 mM EDTA, 1 mM DTT, 100 μg/ml BSA, 0.01% NP40) and cleaved with 0.1 units of Fpg glycosylase. The DNA was lyophilized and resuspended in formamide loading buffer, heated for 3 minutes at 80°C and loaded on 8% sequencing gel in 1XTBE buffer. The gels were dried and exposed overnight on a phosphor imager screen. The screens were scanned on phosphor imager and analyzed by Multi-Gauge (Fuji) software.

### Gel quantification and apparent dissociation constants (K_d_^app^) evaluation

Gel quantifications were performed by integration of rectangles encompassing the cleavage bands of interest. Signal intensities were determined as the relative averaged intensities of the footprinted GG cleavage bands. For a given binding site, these bands are normalized to the “internal standard” bands, belonging to 4-5 other guanines within the respective DNA ladder in the absence of Sox6. The averaged and normalized relative intensities were plotted as a function of the Sox6 concentration together with the mean deviations. To evaluate the apparent dissociation constant (K_d_^app^), the experimental data were fitted mathematically to smoothly decaying curves by least square deviation procedure using either TableCurve 2D 5.01 (81) with arbitrary fitting functions providing R^2^>0.98 and FitStdErr <0.02 or MATLAB R2020B software with biexponential function (fitexp2) providing R^2^>0.97 and RMSE<0.05 (76). The apparent dissociation constant, by analogy with the true K_d_, was determined as the Sox6 concentration corresponding to the 1/2 level of the signal intensity change.

## DATA AVAILABILITY

All the relevant structural models together with the relevant simulation trajectories and their analysis scripts are deposited at GitHub (https://github.com/CSB-KaracaLab/Sox-PTF). Additional data supporting the findings of this study are available from the corresponding authors upon request.

## SUPPLEMENTARY DATA

Supplementary Data are available.

## FUNDING

This work was supported by institutional funding of Izmir Biomedicine and Genome Center; and Centre National de la recherche Scientifique and was benefiting from the 2232 International Fellowship for Outstanding Researchers Program of TÜBİTAK [Project No: 118C354] (the financial support received from TÜBÌTAK does not mean that the content of the publication is approved in a scientific sense by TÜBÌTAK). E.K. acknowledges EMBO Installation Grant (no. 4421), Alexander von Humboldt Foundation Return Fellowship, as well as the Young Investigator Award granted by the Turkish Science Academy.

## ACKNOWLEDGEMENT

All the simulations and analyses were carried out in the TUBITAK’s ULAKBİM High Performance and Grid Computing Center and local HPC resources of IBG.

## CONFLICT OF INTEREST

The authors declare no competing interests.

## Notes

https://github.com/CSB-KaracaLab/Sox-PTF

### Competing Interest Statement

The authors have declared no competing interest.

### Summary of Updates

Main text and SI updated for clarity.

