## Supplementary for "Differential Histone-DNA Interactions Dictate Nucleosome Recognition of the Pioneer Transcription Factor Sox"

**SUPPLEMENTARY FIGURES:**

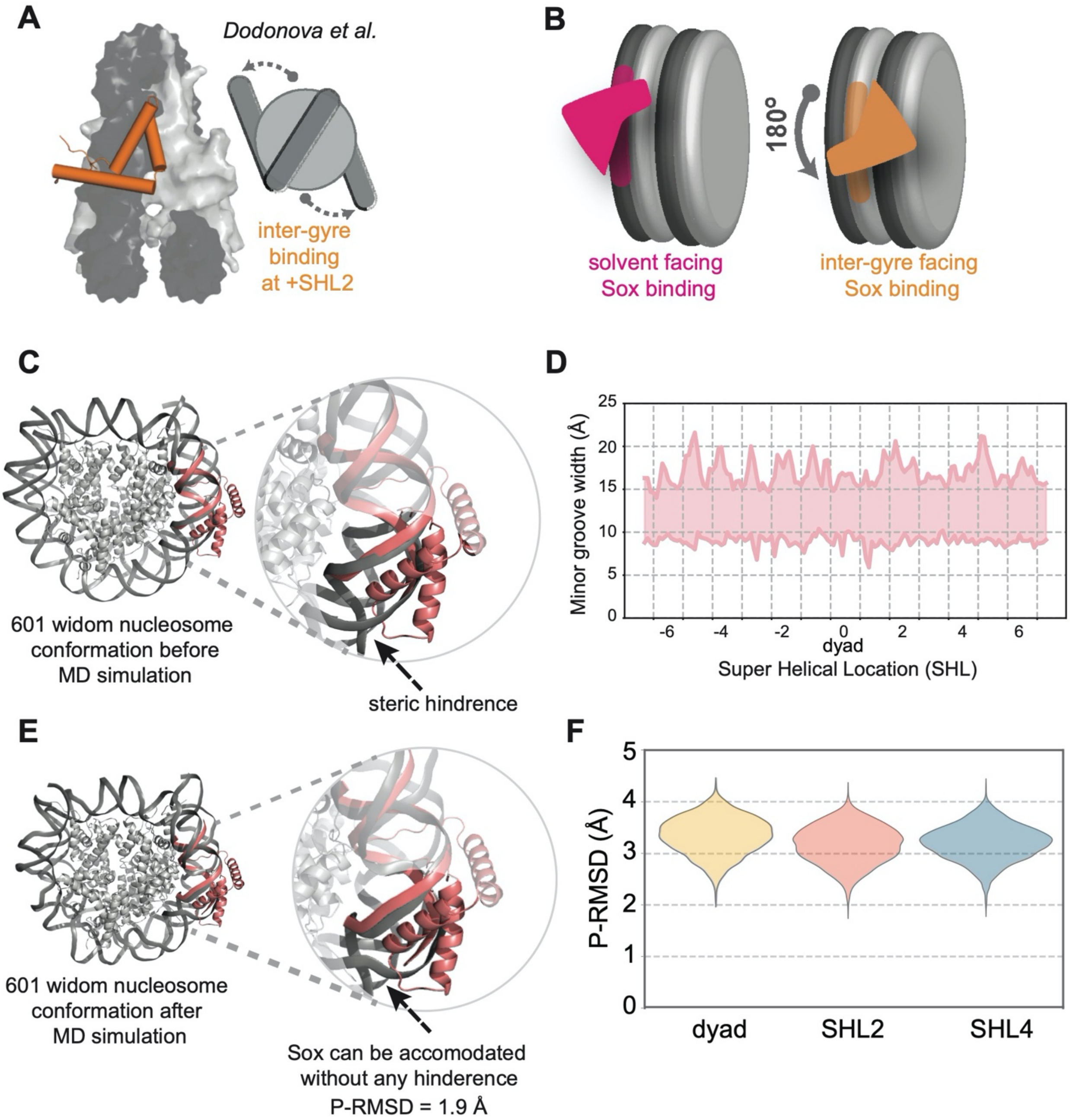


**Supplementary Figure 1. A.** Dodonova *et al.*'s Sox:nucleosome binding mode (pdb id: 6t7b). Sox binding site is located at the inter-gyre facing strand at SHL2. Sox is represented in orange. **B.** The orientation of Sox differs according to the placement of its recognition sequence. If the Sox binding sequence is placed on the positive DNA arm, i.e., on the solvent facing DNA strand (left), Sox should bind to the nucleosome without imposing major structural alterations. **C.** Sox:DNA complex is structurally matched with the Sox recognition sequence, inserted at SHL2. The resulting fitting is highlighted, showing that when free and nucleosomal DNA conformations are matched, Sox (pink) is experiencing serious clashes with nucleosomal DNA. **D.** The minor groove width profile variations of the available nucleosome structures. The signature Sox binding signal requires a peak at 22.5 Å minor groove widening. (The following pdb ids were considered: 1aoi, 1eqz, 1f66, 1id3, 1kx3, 1kx4, 1kx5,

1m18, 1m19, 1m1a, 1p34, 1p3a, 1p3b, 1p3f, 1p3g, 1p3i, 1p3k, 1p3l, 1p3m, 1p3o, 1p3p, 1s32, 1u35, 1zla, 2cv5, 2fj7, 2nqb, 2nzd, 2pyo, 3a6n, 3afa, 3an2, 3av1, 3av2, 3ayw, 3aze, 3azf, 3azg, 3azh, 3azi, 3azj, 3azk, 3azl, 3azm, 3azn, 3b6f, 3b6g, 3c1b, 3kuy, 3kwq, 3kxb, 3lel, 3lja, 3lz0, 3lz1, 3mgp, 3mgq, 3mgr, 3mgs, 3mnn, 3mvd, 3o62, 3reh, 3rei, 3rej, 3rek, 3rel, 3tu4, 3ut9, 3uta, 3utb, 3w96, 3w97, 3w98, 3w99, 3wa9, 3waa, 3wkj, 3wtp, 3x1s, 3x1t, 3x1u, 3x1v, 4jjn, 4kgc, 4kud, 4ld9, 4qlc, 4r8p, 4wu8, 4wu9, 4x23, 4xuj, 4xzq, 4ym5, 4ym6, 4ys3, 4z5t, 4z66, 4zux, 5av5, 5av6, 5av8, 5av9, 5avb, 5avc, 5ay8, 5b0y, 5b0z, 5b1l, 5b1m, 5b24, 5b2i, 5b2j, 5b31, 5b32, 5b33, 5b40, 5cp6, 5cpi, 5cpj, 5cpk, 5dnm, 5dnn, 5e5a, 5f99, 5gse, 5gsu, 5gt0, 5gt3, 5gtc, 5gxq, 5hq2, 5jrg, 5kgf, 5mlu, 5nl0, 5o9g, 5omx, 5ong, 5onw, 5wcu, 5x0x, 5x0y, 5x7x, 5xf3, 5xf4, 5xf5, 5xf6, 5xm0, 5xm1, 5y0c, 5y0d, 5z23, 5z30, 5z3l, 5z3t, 5z3u, 5z3v, 5zbx, 6a5l, 6a5o, 6a5p, 6a5r, 6a5t, 6a5u, 6buz, 6c0w, 6dzt, 6e0c, 6e0p, 6esf, 6esg, 6esh, 6esi, 6fml, 6fq5, 6fq6, 6fq8, 6gej, 6gen, 6hkt, 6hts, 6i84, 6inq, 6ir9, 6iro, 6iy3, 6j4w, 6j4x, 6j4y, 6j4z, 6j50, 6j51, 6j99, 6jm9, 6jma, 6jyl, 6k1p, 6muo, 6mup, 6ne3, 6nj9, 6nn6, 6nog, 6nqa, 6o1d, 6o96, 6om3, 6r1t, 6r1u, 6r25, 6r8y, 6r8z, 6r90, 6r91, 6r92, 6r93, 6r94). **E.** After MD simulations, 601 Widom nucleosome, encompassing the Sox cognate sequence at SHL2 could accommodate Sox (pink) without any clashes. **(F)** The Phosphate Root Mean Square Deviation (P-RMSD) distributions of 601-SHL024 throughout the simulation (N=3000). P-RMSD values are calculated by taking the Sox11-DNA conformation as a reference (pdb id: 6t78). The lowest P-RMSD values are 2.1 Å, 1.9Å and 1.9Å for dyad, SHL2, and SHL4, respectively.

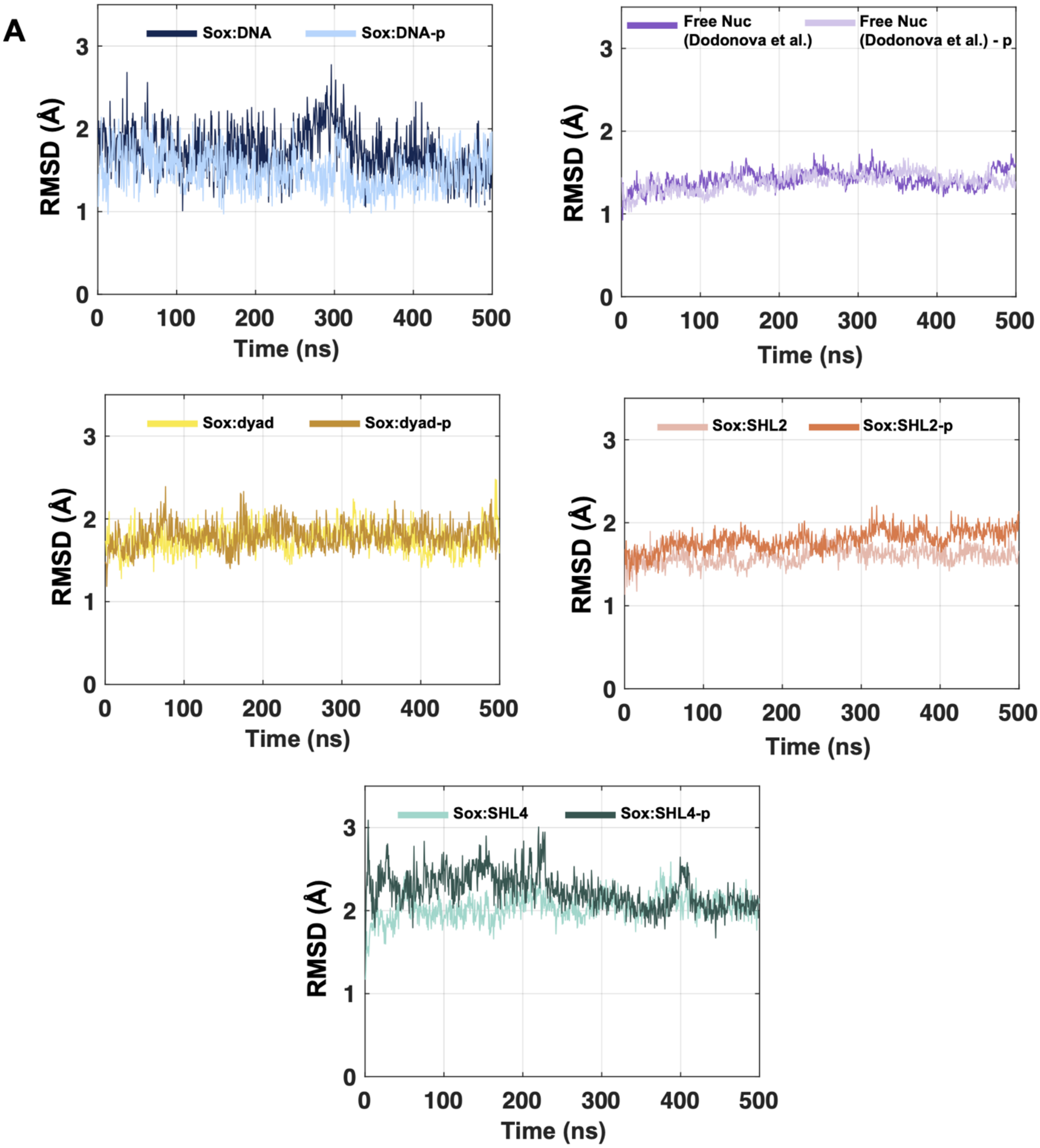
A
Sox:DNA
Sox:DNA-p
Free Nuc (Dodonova et al.)
Free Nuc (Dodonova et al.) - p
Sox:dyad
Sox:dyad-p
Sox:SHL2
Sox:SHL2-p
Sox:SHL4
Sox:SHL4-p
RMSD (Å)
Time (ns)
0
1
2
3
0
100
200
300
400
500

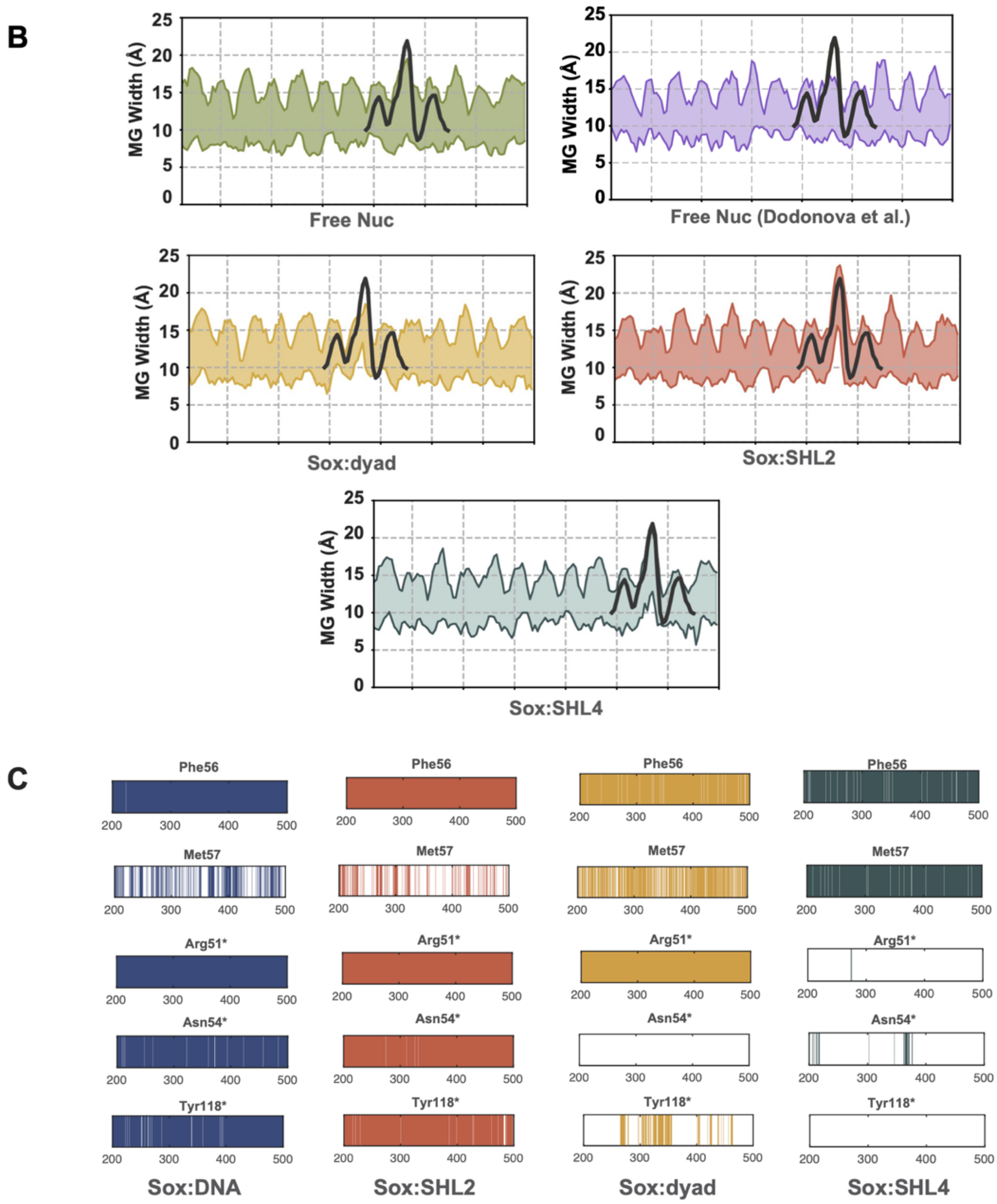


**Supplementary Figure 2. A.** The MD-driven RMSD profiles of free Sox:DNA, free nucleosome determined by Dodonova et al., Sox:dyad, Sox11:SHL2, and Sox11:SHL4 complexes according to their initial conformations. **B.** Minor groove width distributions of each SHL site throughout 601-SHL024 (free nucleosome with SHL0, SHL2, and SHL4 sites mutated), free nucleosome determined by Dodonova et al., Sox11:SHL2, Sox11:dyad and Sox11:SHL4 complex simulations (N=1200 for each complex). The reference profile of Sox SHL2 bound nucleosome structure (pdb id: 6t7b) is shown in dark gray and placed at the relevant SHLs to serve as a comparison. **C.** The Sox11:DNA interaction profiles of the essential Sox amino acids observed in our replica simulations, at free DNA, SHL2, dyad, and SHL4 sites (N=600 for each complex). The amino acid selections follow the ones presented in Figure 1 and 2.

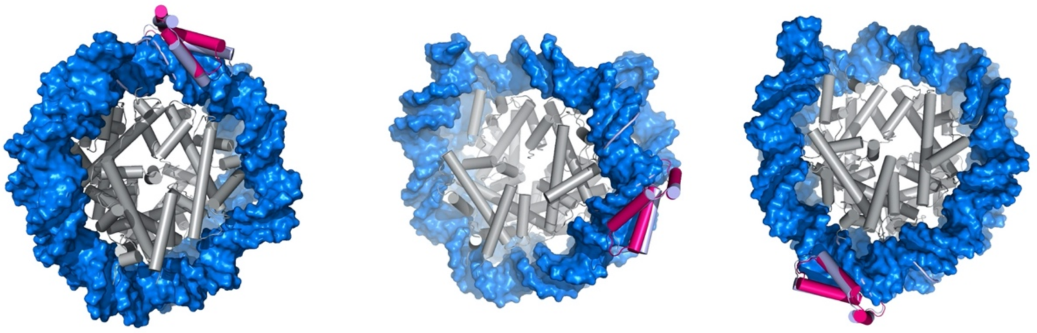


**Supplementary Figure 3.** The lowest P-RMSD Sox6:NCP complexes. The reference Sox:DNA complex (pdb id:6T78) is depicted in lilac.

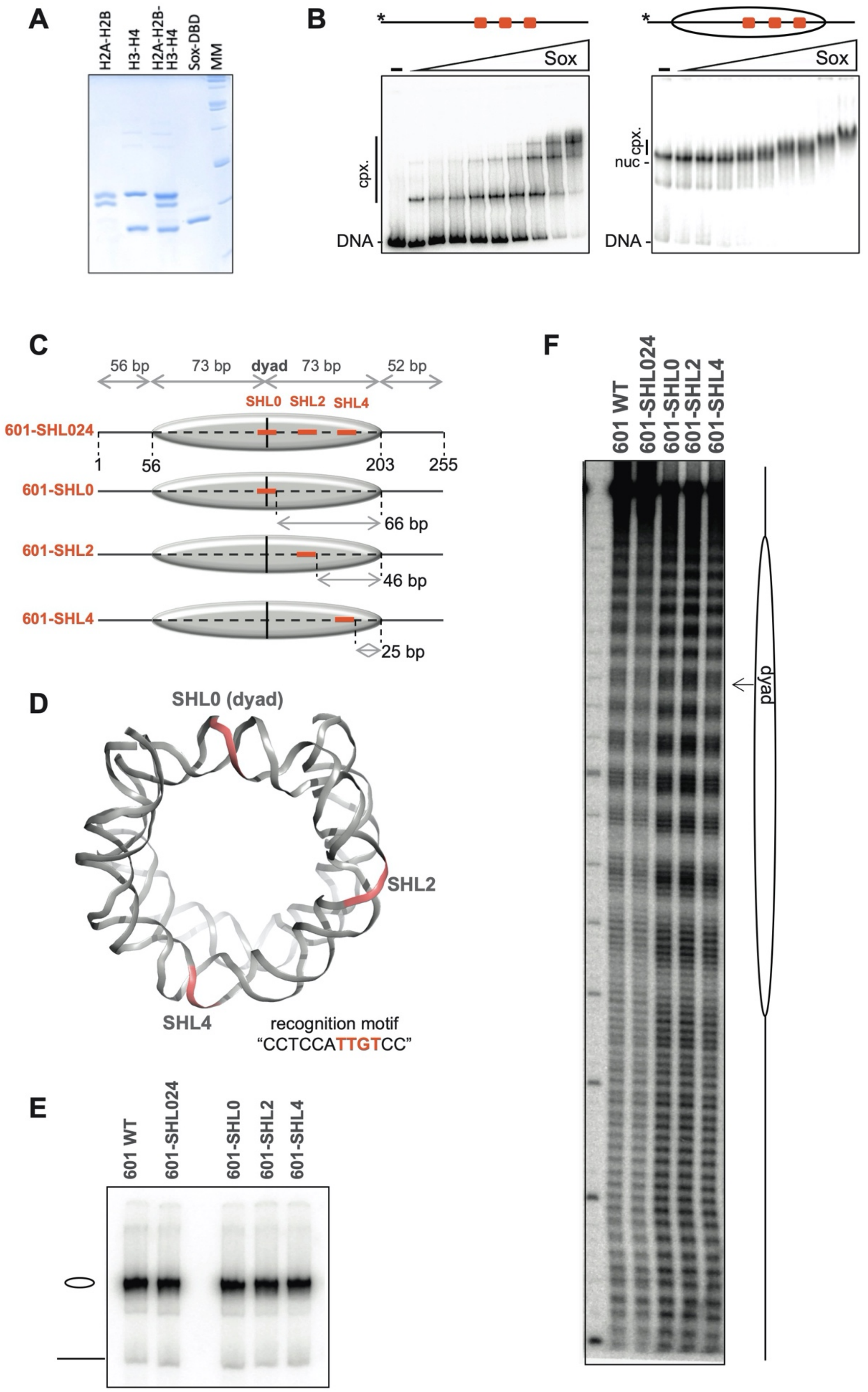


**Supplementary Figure 4. A.** DNA Electrophoretic analysis of the purified recombinant histones, the reconstituted histone octamer and the purified Sox6 HMG-domain. **B.** EMSA showing the binding of Sox6 HMG-domain to naked DNA (601-SHL024) (left) or to nucleosome (601-SHL024) (right). Naked

$^{32}$P-end labeled 601-SHL024 DNA or nucleosomes were incubated with increasing amount of Sox6 HMG-domain and aliquots of the reaction mixtures were run on a native PAGE. For each experimental condition, the concentration of the Sox6 HMG-domain used in the nucleosome binding experiments was ~ 3-fold higher compared to the one used for analyzing the binding to naked DNA. The positions of free DNA, nucleosomes and their complexes with Sox6 HMG-domain are indicated. **C.** Schematics of the reconstituted nucleosomes. The Sox binding site was inserted in the 255 bp 601 DNA fragment at three different locations, namely at the dyad (SHL0), at SHL2 and at SHL4. Bold lines correspond to free DNA arms; oval to core particle region; vertical black line to nucleosome dyad; salmon bold line to Sox binding sites. The numbers refer to the length of DNA in the depicted regions. **D.** 601 nucleosomal DNA sequence (pdb id: 3lz0) is mutated upon inserting Sox binding sequence, *5'-CCTCCA**TTGT**CC-3',* at three different SHLs, namely dyad, SHL2 and SHL4. All the nucleosome inserted Sox binding sites face the solution. The three Sox recognition DNA sequences are colored in salmon. **E.** EMSA of the indicated reconstituted nucleosomes. The positions of free DNA (horizontal black line) and reconstituted nucleosomes (oval) are indicated. **F.** Hydroxyl radical footprinting of the reconstituted nucleosomes. First line, molecular mass marker; Schematics of the nucleosome is shown on the right. The dyad is indicated with an arrow.

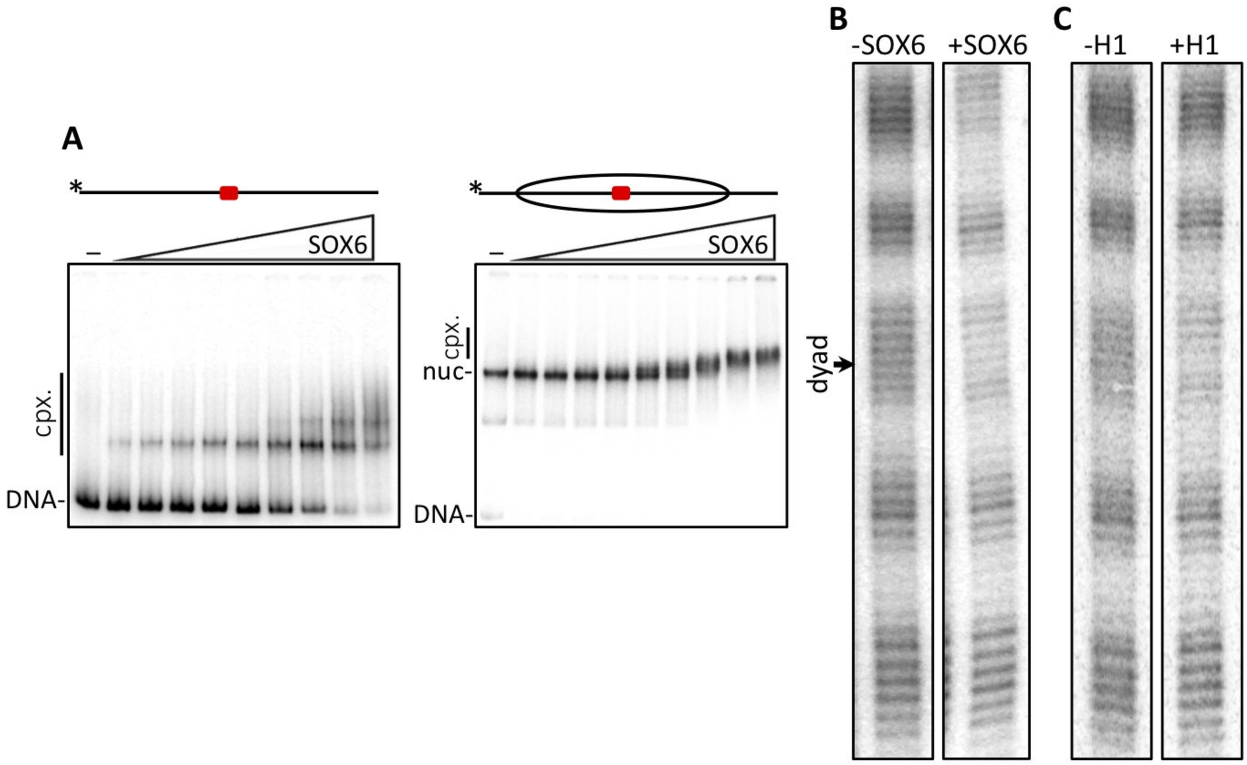


**Supplementary Figure 5. A.** EMSA showing the binding of Sox6 HMG-domain to naked naked (601-SHL0) DNA (left) and (601-SHL0) nucleosome (right). Naked 32P-end labeled 601-SHL024 DNA or nucleosomes were incubated with increasing amount of Sox6 HMG-domain and aliquots of the reaction mixtures were run on a native PAGE. For each experimental condition, the concentration of the Sox6 HMG-domain used in the nucleosome binding experiments was ~ 3-fold higher compared to the one used for analyzing the binding to naked DNA. The positions of free DNA, nucleosomes and their complexes with Sox6 HMG-domain are indicated. **B.** •OH radical DNA cleavage pattern of the region around the dyad of 601 nucleosome with bound (+) and unbound (-) linker Sox6. A clear footprint in the Sox6 bound nucleosome is observed at the dyad. **C.** •OH radical DNA cleavage pattern of the region around the dyad of the 601-SHL024 nucleosome in the absence (-) or in the presence (+) bound H1. The dyad is indicated by an arrow. Note the clear footprint around the dyad of the H1-bound nucleosome.

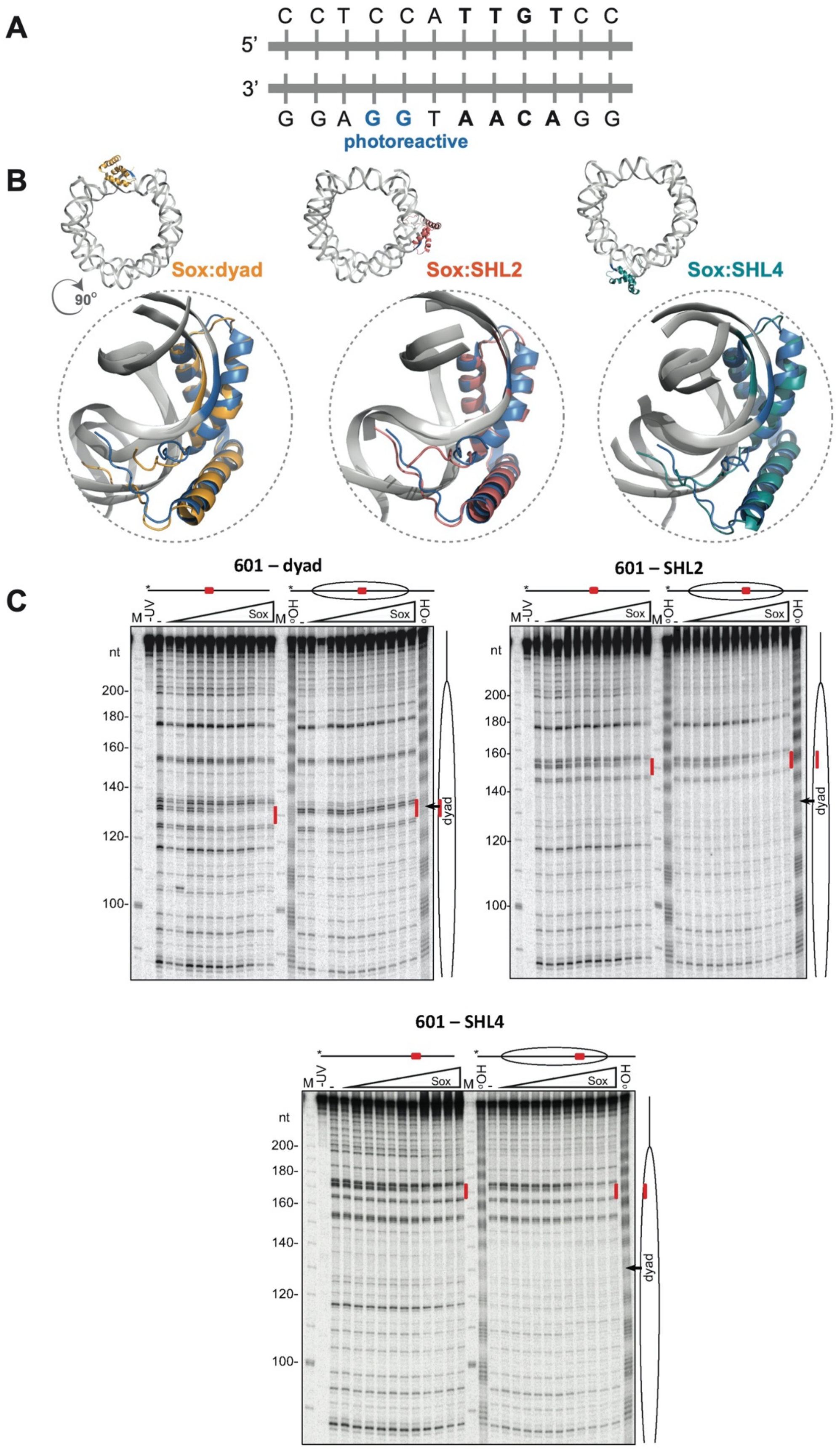


**Supplementary Figure 6. A.** The representative coordinates of Sox cognate binding sequence and photoreactive GG nucleotides. Sox cognate sequence "TTGT" is shown as bold and photoreactive GG nucleotides are depicted in blue. **B.** The representative location of photoreactive GG nucleotides on

SHLs. GG deformation of Sox:nucleosome models are compared with Sox:DNA (pdb id:4y60). Reference is depicted with blue color and Sox:nucleosome at dyad, SHL2 and SHL4 models are presented in orange, pink and green, respectively. **C.** UV laser footprinting patterns of the Sox-DNA and Sox-nucleosome complexes bearing a Sox recognition sequence either at the dyad (left), at SHL2 (right), and at SHL4 (below-middle), respectively. The complexes were irradiated with a single 5 nanoseconds UV laser 266 nm pulse (Epulse,0.1 J/cm$^2$) and DNA was purified from the samples. After treatment of the purified DNA with Fpg glycosylase, the cleaved DNA fragments were separated on 8% sequencing gel and visualized by autoradiography. Red vertical lines and red squares mark the Sox binding sites, M marks the molecular mass, oval represent the schematics of the nucleosome, and the dyad is indicated with an arrow. -UV refers to the control, non-UV irradiated and Fpg treated sample.

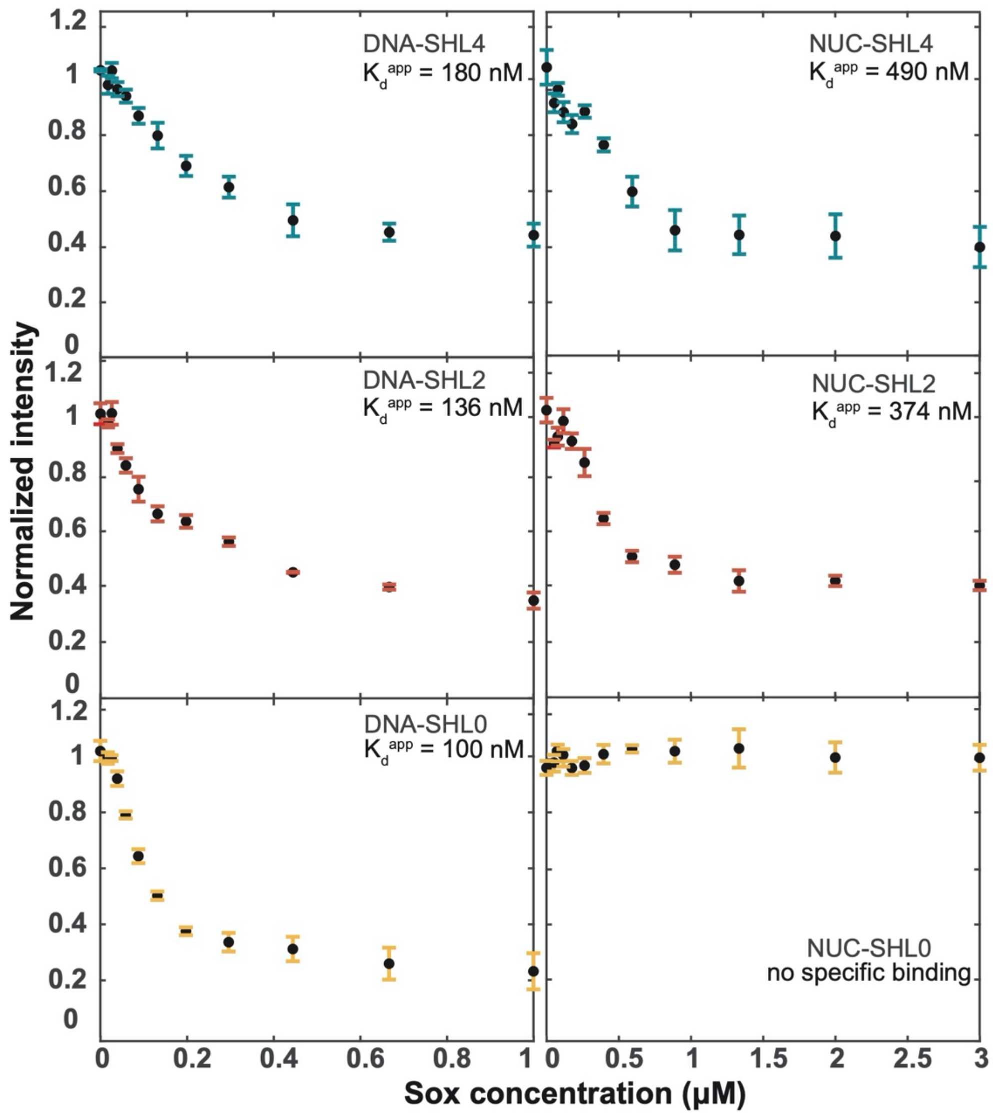


**Supplementary Figure 7.** Sox6 concentration dependences of the footprinting intensity representing the normalized cleavage band intensity of the GG run within the binding site for DNA (left) and Nucleosome (right) bound Sox at SHL4, SHL2 and SHL0 individually.

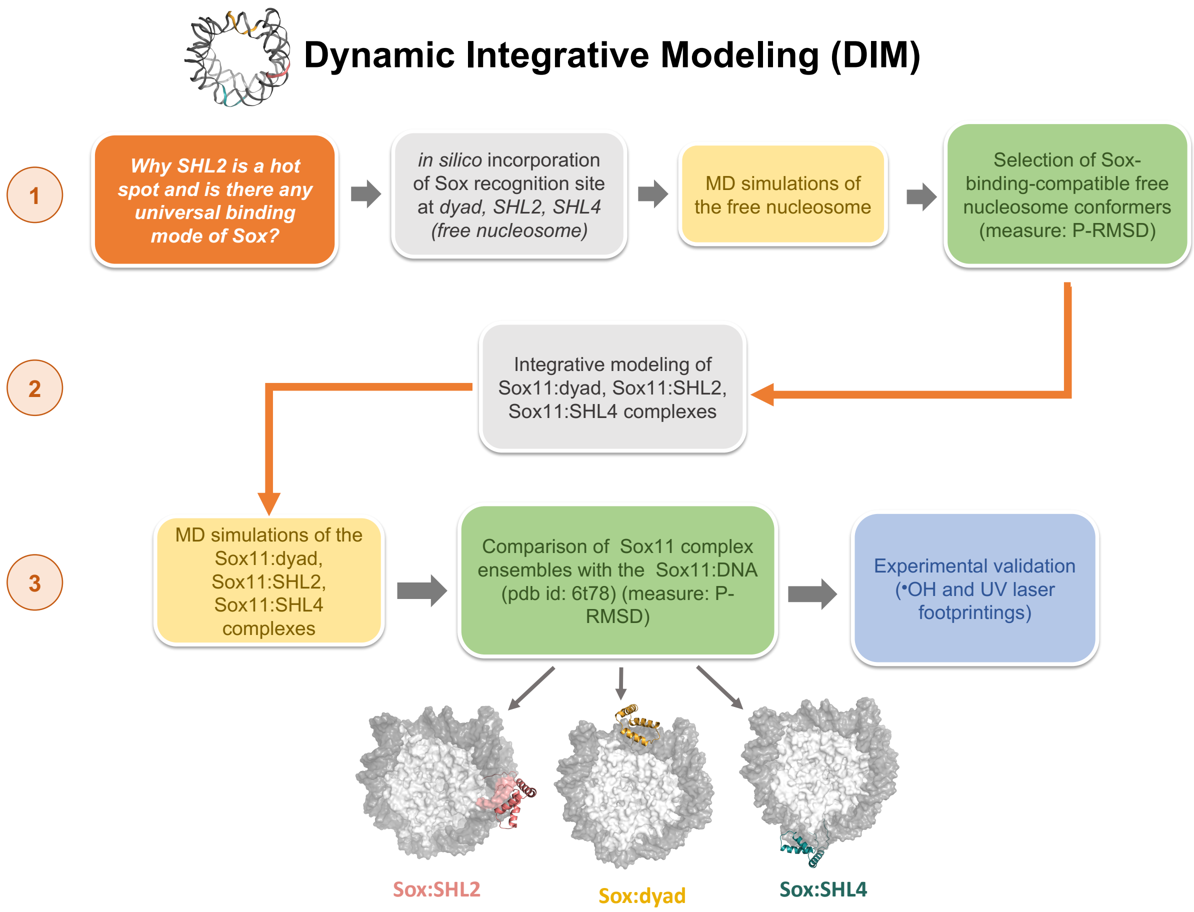


**Supplementary Figure 8. Dynamic Integrative Modeling (DIM) pipeline.** The steps followed during our DIM protocol are indicated within boxes. The orange box is our starting point; yellow and gray boxes refer to MD simulations and modeling steps, respectively; green boxes depict the checkpoints, where we chose a representative structure from the generated conformation pool and blue box represents the experimental validation of our results using ·OH and UV laser footprintings techniques.

**SUPPLEMENTARY TABLES:**

**Supplementary Table 1.** Sox11:DNA interaction percentages of the essential Sox amino acids. Hydrophobic and h-bond base specific interactions are shown in green and purple, respectively. * represents base specific h-bond interactions. The generic (non-specific) interactions are shown in black.

| Sox11 amino acids | DNA:Sox | | Sox:SHL2 | | Sox:dyad | | Sox:SHL4 | |
|---|---|---|---|---|---|---|---|---|
| | Sim1 | Sim2 | Sim1 | Sim2 | Sim1 | Sim2 | Sim1 | Sim2 |
| **Phe56** | **100.0** | **99.8** | **100.0** | **100.0** | **96.8** | **96.3** | **92.0** | **95.8** |
| **Met57** | **54.2** | **39.3** | **42.1** | **25.3** | **68.9** | **74.0** | **87.7** | **97.2** |
| Arg51 | 100.0 | 100.0 | 100.0 | 100.0 | 99.7 | 100.0 | 85.5 | 96.3 |
| **Arg51*** | **100.0** | **100.0** | **100.0** | **100.0** | **70.0** | **100.0** | **77.2** | **0.5** |
| Asn54 | 100.0 | 100.0 | 100.0 | 100.0 | 30.3 | 43.6 | 5.8 | 24.8 |
| **Asn54*** | **99.2** | **97.8** | **100.0** | **99.3** | **0.2** | **0.0** | **0.0** | **4.8** |
| Tyr118 | 99.8 | 100.0 | 99.5 | 99.7 | 91.7 | 69.6 | 17.5 | 28.1 |
| **Tyr118*** | **98.3** | **97.0** | **98.0** | **96.8** | **43.3** | **21.6** | **2.2** | **0.0** |

**Supplementary Table 2.** The Sox11 amino acids that fall within 7Å of the nearest histone protein. The calculations are made on the complexes, fitting to the Sox:DNA binding conformation the best.

| Sox11:SHL2 | Sox11:dyad | Sox11:SHL4 |
|---|---|---|
| ARG121 (C-tail)<br>LYS1211 (C-tail) | SER46 (globular)<br>LYS81 (globular)<br>LYS88 (globular)<br>ARG121 (C-tail)<br>LYS1211 (C-tail) | HIS75(globular)<br>ASN76 (globular)<br>ALA77 (globular)<br>GLU78 (globular)<br>LYS81 (globular)<br>LYS85 (globular)<br>TYR118 (C-tail)<br>PRO120 (C-tail)<br>ARG121 (C-tail)<br>LYS1211 (C-tail) |

**SUPPLEMENTARY MOVIES:**

The link to the Supplementary movies is https://github.com/CSB-KaracaLab/Sox-PTF/tree/main/Movies

SHL2.mov corresponds to the binding mechanism of Sox11 at SHL2; SHL4.mov to Sox at SHL4 and dyad.mov to Sox at dyad, as outlined by our dynamic integrative modeling. The green residues indicate the essential polar Sox amino acids, while the purple ones refer to the hydrophobic ones.
